# AAnet resolves a continuum of spatially-localized cell states to unveil tumor complexity

**DOI:** 10.1101/2024.05.11.593705

**Authors:** Aarthi Venkat, Scott E. Youlten, Beatriz P. San Juan, Carley Purcell, Matthew Amodio, Daniel B. Burkhardt, Andrew Benz, Jeff Holst, Cerys McCool, Annelie Mollbrink, Joakim Lundeberg, David van Dijk, Leonard D. Goldstein, Sarah Kummerfeld, Smita Krishnaswamy, Christine L. Chaffer

## Abstract

Identifying functionally important cell states and structure within a heterogeneous tumor remains a significant biological and computational challenge. Moreover, current clustering or trajectory-based computational models are ill-equipped to address the notion that cancer cells reside along a phenotypic continuum. To address this, we present Archetypal Analysis network (AAnet), a neural network that learns key archetypal cell states within a phenotypic continuum of cell states in single-cell data. Applied to single-cell RNA sequencing data from pre-clinical models and a cohort of 34 clinical breast cancers, AAnet identifies archetypes that resolve distinct biological cell states and processes, including cell proliferation, hypoxia, metabolism and immune interactions. Notably, archetypes identified in primary tumors are recapitulated in matched liver, lung and lymph node metastases, demonstrating that a significant component of intratumoral heterogeneity is driven by cell intrinsic properties. Using spatial transcriptomics as orthogonal validation, AAnet-derived archetypes show discrete spatial organization within tumors, supporting their distinct archetypal biology. We further reveal that ligand:receptor cross-talk between cancer and adjacent stromal cells contributes to intra-archetypal biological mimicry. Finally, we use AAnet archetype identifiers to validate GLUT3 as a critical mediator of a hypoxic cell archetype harboring a cancer stem cell population, which we validate in human triple-negative breast cancer specimens. AAnet is a powerful tool to reveal functional cell states within complex samples from multimodal single-cell data.

## Introduction

Cancer cells can dynamically change their functional state to facilitate survival [8, 15, 31]. This creates a phenotypic continuum of cell states within a tumor that can be viewed as a cell state landscape. In this model, dynamic gene expression changes enable movement about the landscape. Defining the breadth of cell states and their structural organization within a tumor is currently a significant biological and computational challenge, yet is likely to reveal critical opportunities to perturb cancer progression.

The two predominant approaches for characterizing cell state heterogeneity from single-cell transcriptomic data are clustering and trajectory inference [24, 50]. Clustering partitions the cellular state space into discrete cell types, and trajectory inference identifies continuous paths that define a pattern of cellular dynamics. However, when there are no clear delineations between cellular states, nor clear trajectories or lineage structure on the data manifold, neither approach suffices to map the cellular state space. Thus, to define biological similarities and differences within and between these tumors, there is a need for a method that can dissect cellular heterogeneity at single-cell resolution while maintaining continuous variation along the cell state continuum. For this purpose, we turn to archetypal analysis, a factor analysis technique first introduced by Cutler and Breiman [11]. Archetypal analysis first extracts factors that represent the “archetypes”, or extreme states, of a dataset. Then, all datapoints can be described as a convex combination of archetypes. In other words, archetypal analysis models the data as a simplex, where the extreme points are corners and other points are on the faces or are internal to the simplex (Figure 1). However, viewing this technique geometrically makes it clear that most datasets would not be accurately described by such a simplex in the ambient data space.

**Figure 1.**
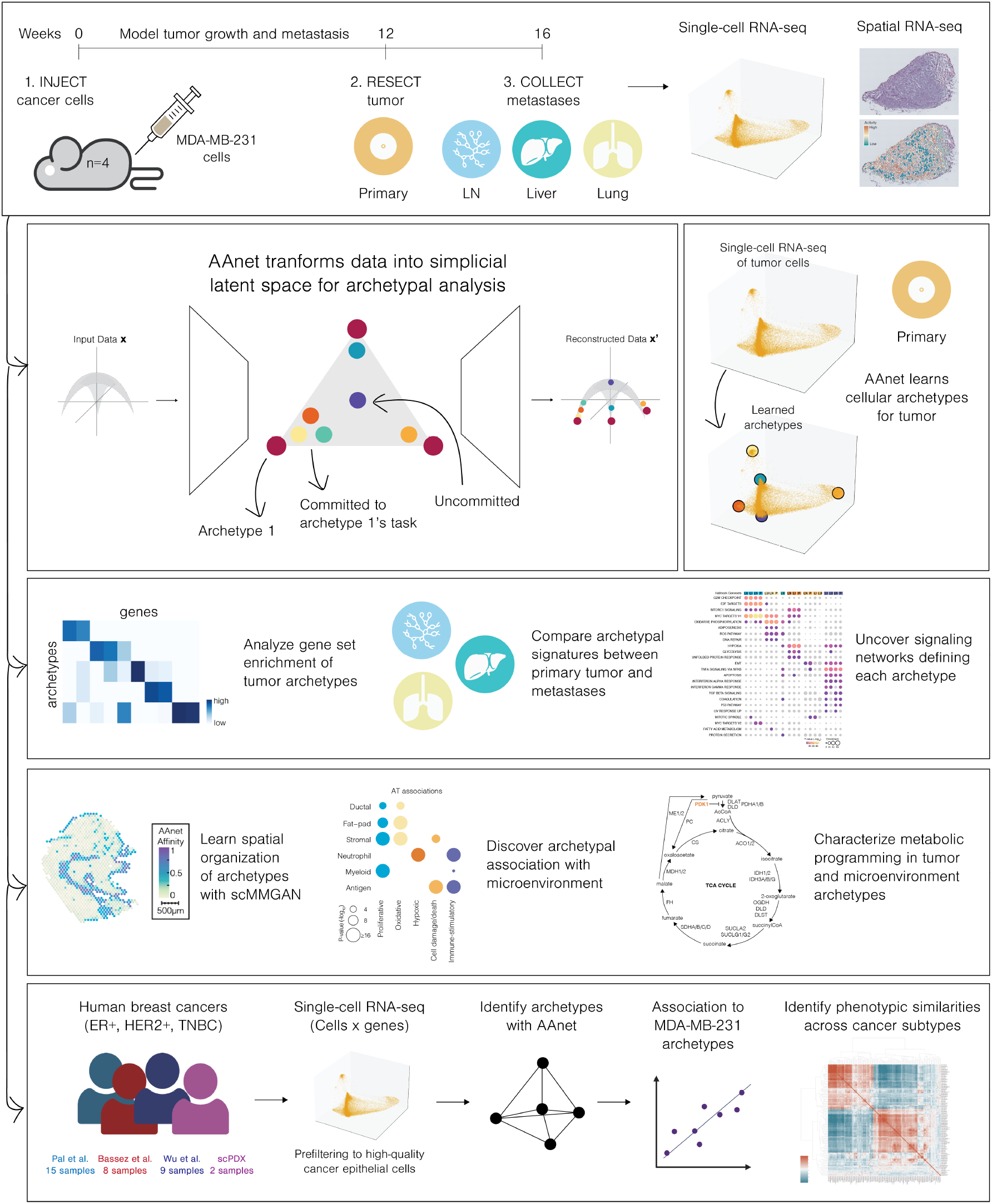
Overview of experimental design, AAnet, and downstream analyses.

This motivated the development of our approach — Archetypal Analysis network (AAnet) — which performs archetypal analysis by *learning* a simplicial representation of the data. AAnet is implemented as an autoencoder, i.e. a neural network that learns meaningful representations of the given data, but is regularized to perform archetypal analysis. Instead of fitting a simplex on the data space, the AAnet encoder learns the optimal transformation from the data space to a latent space bound by a simplex, and the decoder learns the transformation back to the data space. This means that AAnet learns archetypes and a representation of each datapoint as convex combination of archetypes through nonlinear dimensionality reduction. This adds flexibility to archetypal detection while preserving data geometry.

Triple-negative breast cancer (TNBC) is a particularly heterogeneous subtype of breast cancer as it is an amalgamation of all “other” breast cancers that cannot be classified as ER+, PR+, or HER2+. With a lack of specific markers to characterize TNBC, there are consequently a paucity of effective targeted therapies to treat it. Here, we use AAnet to deconvolute TNBC heterogeneity into biologically interpretable archetypes. Using novel single-cell RNA sequencing (scRNAseq) and spatial transcriptomics datasets modeling tumor formation and metastasis *in vivo*, AAnet identifies five archetypes in primary tumors, demonstrating they are reproducibly defined by distinct cancer hallmarks. These include a proliferative archetype associated with cell cycle progression; an oxidative archetype associated with oxidative phosphorylation, ROS production, and adipogenesis; a hypoxic archetype enriched for enzymes associated with oxygen-independent glycolysis; a cell damage/death archetype that captures variation introduced by technical factors; and an immune-stimulatory archetype with enriched expression of HLA genes and cytokines. We validate these archetypes by showing they are recapitulated in metastases, are spatially organized, and colocalize with distinct microenvironmental cells types and metabolic niches. Moreover, in a cohort of 34 human TNBC samples, AAnet reveals significant archetypal heterogeneity between patients. Notably, we identify a subset of patients defined by the hypoxic archetype favoring residence in a cancer stem cell niche, and we validate GLUT3 as a critical regulator of that archetypal cell state. These findings highlight the powerful ability of AAnet to define biological function and organization within cancer, potentially aiding in classifying patients according to biological similarities within and across patient samples. Furthermore, AAnet defines core transcriptional programs driving distinct archetypes, thereby suggesting potential therapeutic opportunities to target them.

## Results

### Defining *in vivo* cellular heterogeneity in scRNAseq data of triple-negative breast cancer Overview of AAnet

We, and others, have shown that cancer cells reside along a phenotypic continuum [8]. The ability to identify critical cell states along the continuum will garner insight into the molecular programs enabling cell state adaptation, thereby facilitating therapeutic strategies to prevent it. For characterizing cell state heterogeneity from scRNAseq data, clustering and trajectory inference are considered a standard part of single-cell workflows and best practices [30]. However, we show that when the data lies on a continuum without latent cluster structure (e.g., discrete cell types) or latent lineage structure (e.g., development axis), these approaches lack concordance across methods and are limited in their ability to meaningfully characterize the cellular state space (Figure 2a, Supplementary Figure 1a-c).

**Figure 2.**
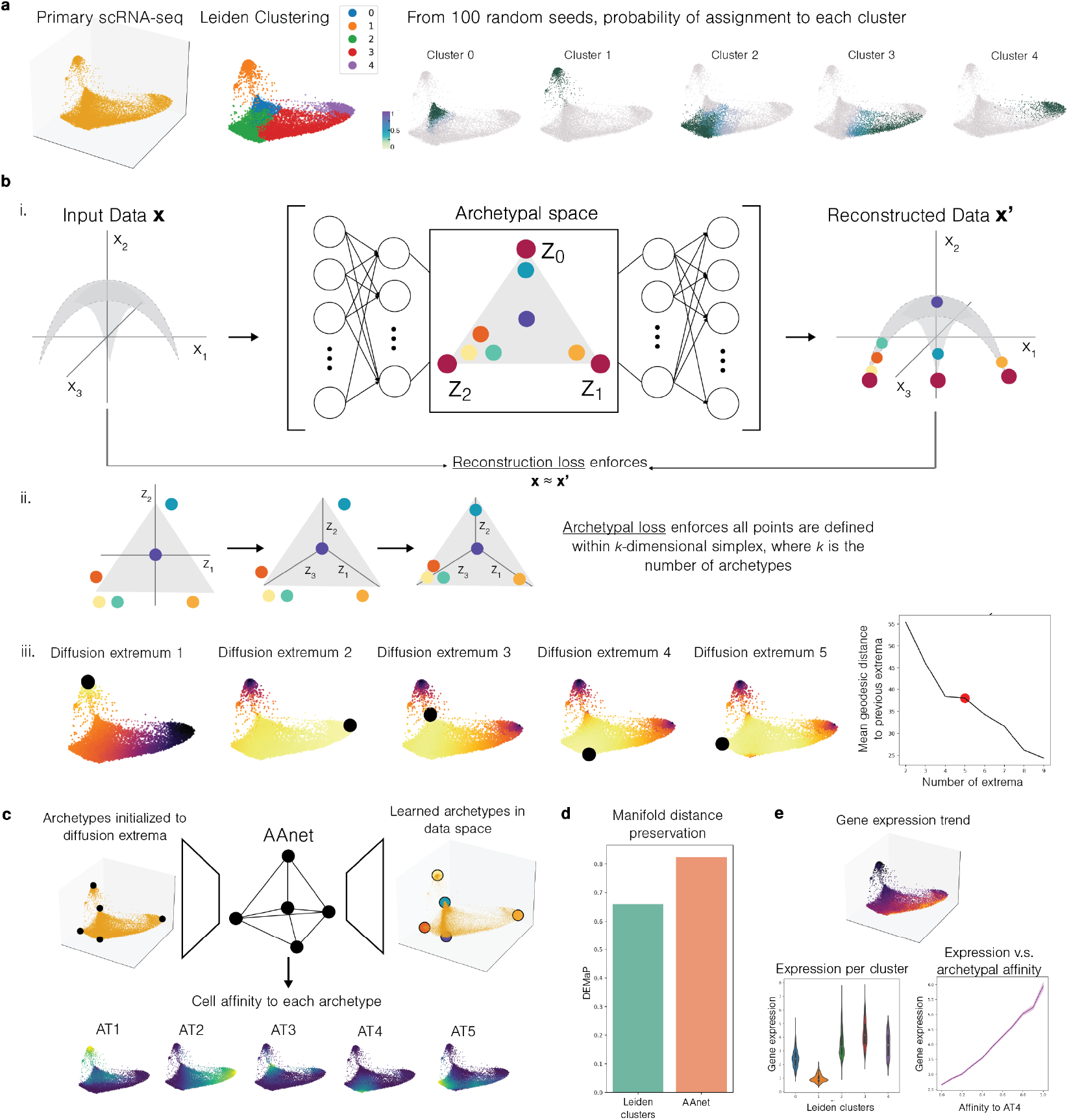
Overview of AAnet architecture. (**a**) Primary tissue visualization is a continuum of cells. Clustering the data with standard clustering tools 100 times (with no parameters changed) results in shifting boundaries between adjacent clusters. (**b**) i. AAnet learns transformation of data into a latent space shaped as a simplex. ii. Archetypal loss enforces that data lies in a latent space shaped like an *k*-dimensional simplex. iii. Diffusion extrema loss infers the extrema from the data geometry. The diffusion extrema also inform the number of archetypes for downstream analysis. (**c**) AAnet is initialized with diffusion extrema and learns affinity to each archetype and decoding of archetypes the data space. (**d**) Manifold distance preservation score (DEMaP) [32] of cluster representation versus AAnet representation. (**e**) AAnet latent space can be used to characterize continuous gene trends not easily characterizable with clustering.

Archetypal analysis provides a framework to identify “archetypes”, or extreme states, in a dataset and characterize the datapoints as a convex combination of archetypes [2, 11, 21, 38]. Currently, the biggest challenge of archetypal analysis is identifying the correct and relevant archetypes. Linear archetypal analysis methods, such as Principal Convex Hull Analysis (PCHA), fit a linear simplex onto the data in the ambient space [11] but fail to correctly identify the extrema when the extrema in the ambient space do not conform to the data geometry. These methods thus prove inflexible to more complex datasets, such as scRNAseq data.

To this end, we developed AAnet, a neural network for nonlinear archetypal analysis (Methods). AAnet learns a low-dimensional latent representation of the data as a regular simplex (Figure 2b i). This is achieved by regularizing the encoding layer of the neural network to encode points as convex or barycentric coordinates based on the archetypal points (Figure 2b ii). The autoencoder-style (i.e. encoder-decoder) architecture and the archetypal regularization together ensure that the model learns an accurate transformation to a simplex representation of the data, and it can decode data from the simplicial space to generate new data. In order to add further robustness to noise and accuracy to the model, we developed an approach (Methods) to identify extreme points based on the underlying data geometry, termed *diffusion extrema* (Figure 2b iii). We then use geodesic distances between diffusion extrema to choose the number of archetypes *k*, and we initialize the archetypes to the first *k* diffusion extrema at the beginning of training.

Once the model is trained, the simplicial latent representation can be used for exploration of the dataset. The vertices in the latent space, encoded by standard basis vectors, can be decoded to the archetypes in the gene space, enabling characterization and comparison of their expression profiles. Furthermore, the archetypal space coordinates provide an interpretable measure of each cell’s affinity to each archetype (Figure 2c, Supplementary Figure 1g). Importantly, these archetypal affinities retain more information about cellular relationships than clustering while maintaining the interpretability of cell types. We show that the simplicial latent representation better preserves geodesic distances [32] than the cluster representation (Figure 2d), suggesting it could prove useful in tasks that depend on cluster annotations (e.g. [17, 27]). Finally, we can also represent signals, including gene expression, with respect to archetypal affinities, which allows characterization of continuous signals not possible with cluster-based enrichment analysis (Figure 2e).

### Comparison of AAnet to other approaches on simulated data

To compare AAnet with existing approaches for characterizing cell state heterogeneity and archetypal analysis, we generated a nonlinearly-transformed tetrahedron, or a simplex with four vertices that are “ground truth” archetypes. We also simulated a nonlinear signal based on the true archetypal affinity to vertex four (Supplementary Figure 1a). Clustering (Supplementary Figure 1b) and trajectory inference (Supplementary Figure 1c) approaches show disagreement in cluster and pseudotime assignments respectively and fail to capture the underlying relationship between the simulated signal and vertex four. Additionally, existing archetypal analysis methods cannot correctly infer the vertices of the tetrahedron and show worsening performance as we increase the nonlinearity of the tetrahedron transformation, suggesting that the linearity of these approaches is their major limitation (Supplementary Figure 1d).

By contrast, AAnet is able to infer the true vertices with nearly perfect performance at all levels of nonlinearity (*γ*_*extrema*_ = 5, Supplementary Figure 1e). Without the diffusion extrema loss (*γ*_*extrema*_ = 0), AAnet shows better average performance than existing methods, though lacks robustness at very high degrees of nonlinearity (Supplementary Figure 1e). The approach to identify the number of archetypes based on diffusion extrema correctly identifies four vertices (Supplementary Figure 1f). Finally, AAnet captures interpretable archetypal affinities, and plotting the simulated signal against the inferred vertex 4 archetypal affinity shows AAnet is able to recapitulate the sinusoidal relationship (Supplementary Figure 1g).

### Validation of AAnet on an antigen-specific CD8+ T cell dataset

To validate our method on real biological data, we leveraged a published single-cell dataset of tumor-specific CD8+ T cells [10]. (Supplementary Figure 2a).

Using a mouse model of lung adenocarcinoma designed to express neoantigens, Connolly et al. identified a reservoir of antigen-specific stem-like T cells in the tumor-draining lymph node (dLN), which then migrated to the site of the tumor to terminally differentiate. Tumor-specific CD8+ T cells from early (8 weeks p.i.) and late (17 weeks p.i.) dLNs (top left), early and late tumors (top right), early dLNs and early tumors (bottom left), and late dLNs and late tumors (bottom right) (Supplementary Figure 2a) were co-embedded with PHATE [32]. “Flags” (blue, white, green) were hand-annotated based on a continuum of expression of key immune-related marker genes in distinct regions of the single-cell embeddings. Given the utility of these flags to characterize the heterogeneity of the state space without demarcating boundaries between points, we refer to these flags as archetypes.

The white archetype was characterized by a naive CD8+ T cell signature based on high expression of *Sell, Lef1*, and *Ccr7*. The green archetype was characterized by a stem-like signature, defined by high expression of *Tcf7, Xcl1, Slamf6*. The blue archetype was characterized by an exhausted signature, with high expression of *Pdcd1, Havcr2*, and *Cd101*.

Here, we ran AAnet separately on each of the paired embeddings to determine if it could recapitulate these archetypes (Supplementary Figure 2b). First, we compared expression of each marker gene to characterize the archetypes. This resulted in annotation of one archetype in each of the paired embeddings as a naive archetype, one archetype in three out of four paired embeddings as stem-like, and one archetype in three out of four paired embeddings as exhausted (Supplementary Figure 2c). Three archetypes (LN AT2, Week 8 AT2, and Lung AT3) did not strongly express any of the markers. These correspond to the hand-annotated exhausted archetype in the lymph node, an uncharacterized part of the manifold, and the hand-annotated stem-like archetype in the lung, respectively. The authors note in the text that, while annotated, there was no prominent population of exhausted T cells in the lymph node and no prominent population of stem-like T cells in the lung. The uncharacterized archetype does not correspond to these three cell state extremes, possibly corresponding to an uncovered cell state.

With the archetypes expressing the markers of interest, we computed pairwise cosine similarity across all measured genes (not only the markers of interest, as in the original work). This showed clear clustering corresponding to the naive, exhausted, and stem-like archetypes (Supplementary Figure 2d). This bolsters the finding in the paper by suggesting that these cell states share broader transcriptomic similarity not limited to the nine markers known to be associated with CD8+ T cell states.

Finally, to highlight the utility of the latent space learned by AAnet, we plotted the marker expression versus the latent space score for each archetype in the week 17 embedding (Supplementary Figure 2e). In all cases, the corresponding marker genes are upregulated, and the other marker genes are downregulated. Furthermore, we see non-linear dynamics of gene patterns corresponding to distance in the latent space, adding an additional layer of information through which to interpret the results.

Together, this analysis corroborates the use of AAnet for characterizing datasets with continuous and nonlinear structure with respect to archetypes. Furthermore, it validates the ability to compare archetypes across datasets to identify unified signatures of response.

### AAnet identifies five major archetypal expression states in primary tumor of triple-negative breast cancer model

Having demonstrated the power of AAnet to deconstruct single cell data into meaningful archetypes (ATs), we sought to use AAnet to address a critical question in cancer biology — how does the cell state landscape change across primary and metastatic tumors? Resolving this question may lead to new strategies to prevent non-genetic adaptation that facilitates cancer progression.

To answer this, we generated a new scRNAseq dataset using an *in vivo* model of triple-negative breast cancer. Highly metastatic MDA-MB-231 breast cancer cells were injected into the mammary fat pad of NSG mice and left to grow (6-8week, Female, n=4; Figure 1). At 12 weeks, mice underwent survival surgery to remove primary tumors and allow metastases to develop further. Primary tumors were dissociated into single cells, sorted by flow cytometry to capture the human cancer cells only (CD298+ cells [26]) and immediately captured for scRNAseq. After an additional 4 weeks *in vivo*, lung, liver and lymph nodes were harvested and CD298+ human tumor cells were sorted and captured for scRNAseq. This hybrid human-mouse model has a number of important strengths that are well suited for this study: 1) the confident delineation of tumor cells from the surrounding microenvironment 2) the capture of matched tumors and metastases and 3) a homogeneous starting population to control for cell intrinsic differences such as genetic clonality. With this model, we used AAnet to deconvolute the archetypes underlying heterogeneity in primary tumors. A total of 28,478 cells were analyzed from four primary-tumors (5,118-8,163 cells) after quality control (Methods).

We first examined the archetypes contributing to cellular heterogeneity within primary tumors (Figure 3a). Each primary tumor was analyzed with AAnet individually (Tumor 1-4), as well as for all primary tumors combined (All). As hypothesized, each dataset showed a continuum of cellular expression states with multiple extrema, rather than discrete clusters or unidirectional trajectories (Figure 3b, Supplementary Figure 3, Supplementary Figure 4a). Five archetypes were identified in each dataset, representing distinct biological roles (Figure 3b, Supplementary Figure 3). To elucidate the biology underlying these archetypes, marker genes were defined and used to identify hallmark genesets upregulated in each archetype and conserved between replicates (FDR < 0.05) (Figure 3d, Supplementary Table 1, Methods). These genes and genesets summarize the biology of each archetype as follows:

**Figure 3.**
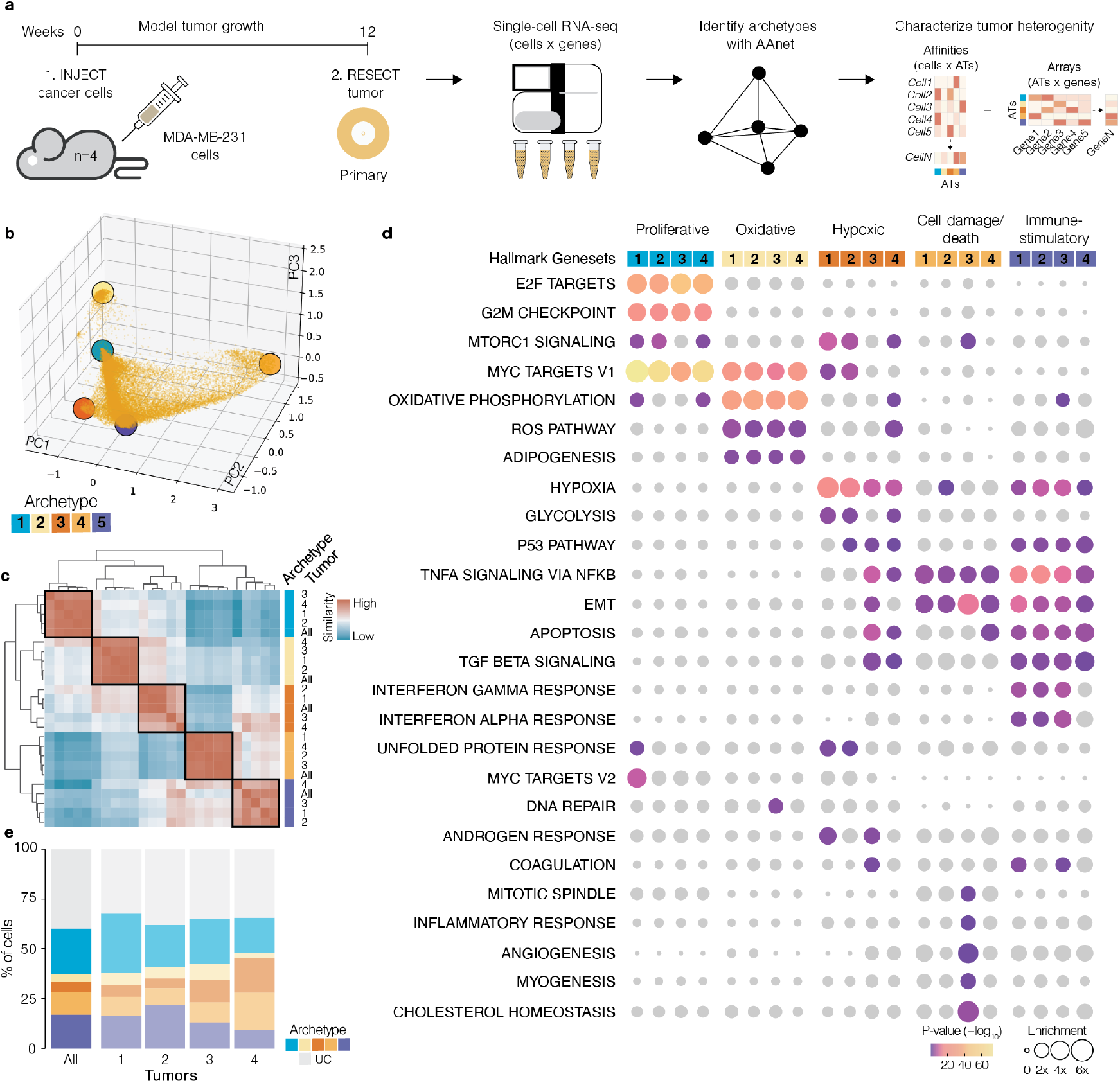
(**a**) Experimental approach used to identify tumor archetypes in TNBC with AAnet. (**b**) Embedding of TNBC tumor cells (all tumors combined) with archetypes indicated by colored circles. (**c**) Heatmap of cosine similarity between archetypal expression vectors determined in each individual tumor (numbered) and all tumors combined (all). Orthologous archetypes between samples are indicated with colors. (**d**) Enriched HALLMARK genesets associated with each archetype in each tumor (colors indicate AT, numbers indicate tumor). P-value is false discovery rate corrected, gene sets where p > 0.05 are shown in gray. Enrichment represents log2 fold-change. (**e**) The percentage of cells in each sample committed to each archetype (colored) or uncommitted to an archetype (gray).

### Proliferative archetype (blue)

This archetype is enriched for hallmarks of cell proliferation (Hallmark genesets G2M checkpoint, E2F targets) and growth (MYC Targets V1/2, mTORC1 signaling). The top markers of this cluster include CDC20, CDK1 and CDK4, key regulators of phase transitions during the cell cycle [47]. Concomitantly, analysis of cell cycle in the cells most strongly associated with this archetype revealed that >95% were in either the S or G2M phase (Supplementary Figure 5a-b).

### Oxidative/adipogenic archetype (yellow)

This archetype is associated with hallmarks of oxidative metabolism and stress. Oxidative phosphorylation (OXPHOS) is the most strongly enriched hallmark geneset across all replicates, driven by electron transport chain components which couple ATP-synthesis to oxygen availability in the mitochondria (Supplementary Table 1, [5]). MYC targets are also overrepresented, including many nuclear genes involved in mitogenesis. Genes in the reactive oxygen species (ROS) pathway are among the top markers of this archetype, including the 4/6 peroxiredoxin family of antioxidant enzymes (PRDX1/2/4/6), regulated by cancer cells in response to oxidative stress [36]. Genes involved in adipogenesis are also significantly overrepresented in all replicates, suggesting fatty acid synthesis may be important for this archetype.

### Hypoxic archetype (orange)

This archetype was significantly associated with the hypoxia hallmark in all replicates. While the canonical regulator of hypoxia HIF1A (hypoxia inducible factor 1) was not among the markers of this archetype, likely because it is degraded post-translationally in the presence of oxygen, CITED2, a HIF1A induced regulator of hypoxia known to promote both breast cancer, was one of the top markers associated with this archetype (Supplementary Table 1, [16]). Enrichment of this geneset was also driven by glycolytic enzymes among marker genes (the glycolysis hallmark enriched in 3/4 replicates). These included class 1 glucose transporters SLC2A1, SLC2A3 as well as genes linked to oxygen-independent energy production in cancer ENO2, HK2, LDHA, GAPDH [1]. Ribosomal subunits also featured prominantly among marker genes, suggesting a relationship between ribosome biogenesis and hypoxia.

### Cell damage/death archetype (amber)

The top markers for this archetype were genes encoded on the mitochondrial genome, for which enrichment is associated with cell damage or death (Supplementary Table 1, [30]). Indeed, analysis of the cells most strongly influenced by this archetype showed high expression of mitochondrially encoded components of the electron transport chain (ETC), yet limited expression of somatically encoded ETC components (Supplementary Figure 6). In addition to mitochondrial genes, TNF signaling via NF-κB and the epithelial-to-mesenchymal transition (EMT) hallmark genesets, both associated with cell stress, were also enriched. This associates this archetype with damaged and dying cells, potentially arising from technical variables.

### Immune-stimulatory archetype (purple)

Immune stimulatory proteins were among the top markers of this archetype, including CXCL1, IFITM2/3, BST2, HLA-A/B/C, B2M and ICAM1 (Supplementary Table 1). Concordantly, Hallmark analysis showed enriched genesets related to immune signaling (IFN-γ, IFN-α, TNF-α, and TGF-β pathways), and apoptosis (Apoptosis, p53). Analysis of cell cycle among the cells most strongly influenced by this archetype showed 96% percent of cells in G1, perhaps suggestive of a G1 arrest (Supplementary Figure 5a-b). These analyses indicate this archetype describes an apoptotic, immune-stimulatory expression pattern in tumor cells.

Archetypes were highly conserved between primary tumors (Figure 3c). All archetypes had an orthologous archetype in each individual primary tumor and the combined dataset, with a mean cosine similarity 0.77-0.97. In contrast, individual archetypes showed modest pairwise cosine similarity (between −0.15 and −0.2), consistent with their classification of representing distinct cell states.

Having deconvoluted tumor cell heterogeneity into five biologically meaningful archetypes, we then determined the association of each cell with each archetype using AAnet. AAnet encodes a representation of each cell based on its relative association to each archetype, where the coordinates of each cell are non-negative and sum to one. We term this association “archetypal affinity” and define cells with affinity for one archetype greater than the sum of all other archetypal affinities (i.e. affinity > 0.5) as “committed”. Cells that do not surpass this affinity threshold for any archetype can be considered “uncommitted”. Using archetypal commitment to analyze the cellular composition of primary tumors, the abundance of cells committed to an archetype was consistently above 60% in all tumors (62%-67.8%), yet the abundance of committed cells varied between archetypes (Figure 3e). The proliferative archetype was the most abundant archetype in the combined analysis and in two of four individual tumors, and the immune-stimulatory archetype was the second most abundant archetype in the combined analysis and in two of four individual tumors. The oxidative/adipogenic archetype represented a minor fraction of tumor cells across all samples, and the hypoxic archetype was among the most variable. Together, these analyses show that AAnet deconvolutes cancer cell heterogeneity into biologically meaningful archetypes that are reproducibly detected in discrete tumors.

### AAnet reveals conserved and *de novo* heterogeneity across distinct metastatic sites

Next, we used AAnet to identify archetypes in metastases. scRNAseq was performed on matched lymph node (LN), liver, and lung metastases that were collected four weeks after resection of the primary tumor (Figure 4a-c). A total of 42,250 metastatic cells were analyzed from LN (n = 4,6604-19,224 cells per tumor), 42,647 cells from liver (n = 4,7136-18,671 cells per tumor), and 17,687 cells from the lung (n = 3,5599-6,379 cells per tumor) after removal of one outlier lung sample during QC (Methods, Supplementary Figure 7a). We combined data from metastases per tissue separately and defined archetypes in each site using AAnet.

**Figure 4.**
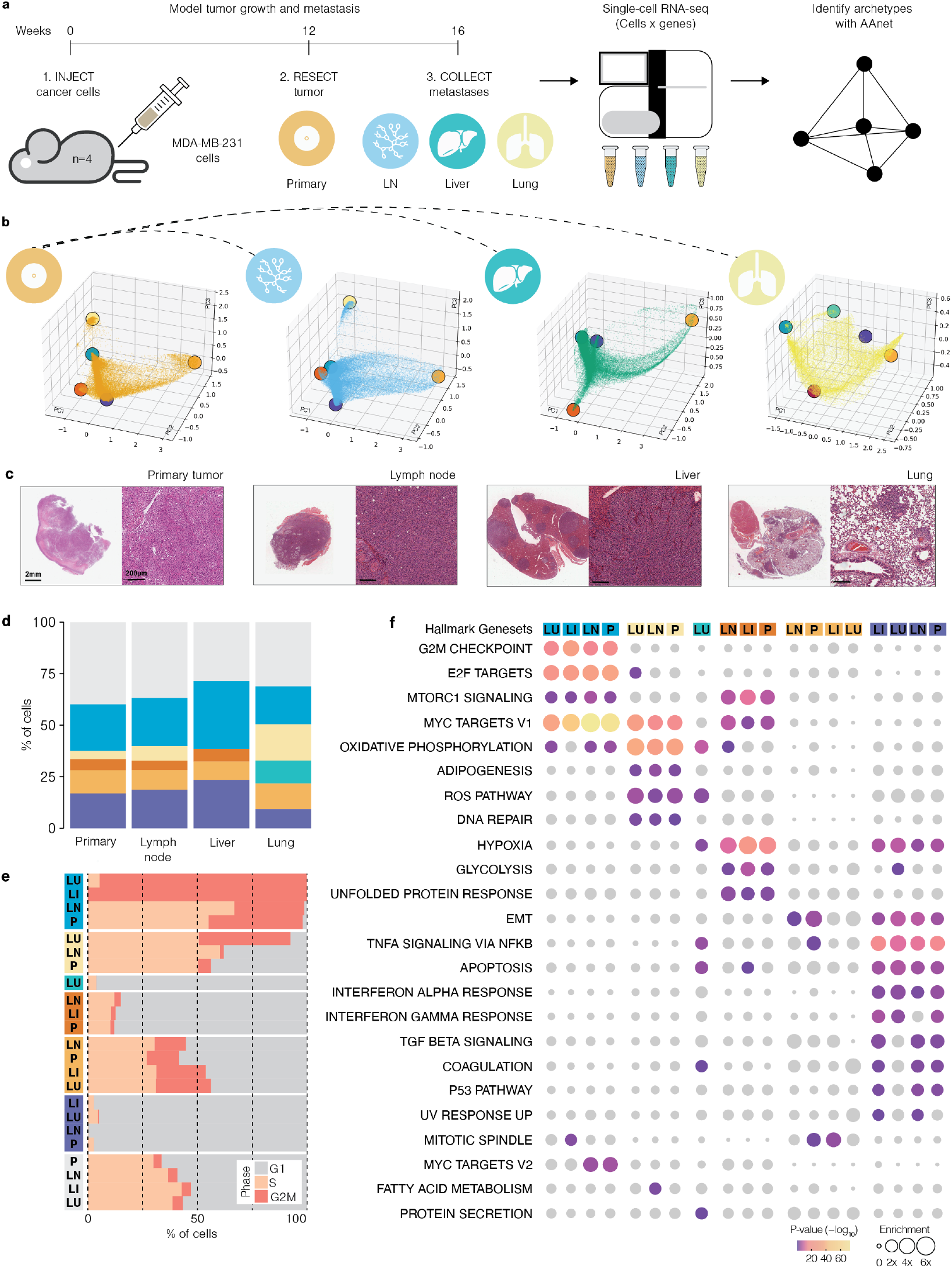
(**a**) Approach to define tumor archetypes in TNBC metastases from the lymph node (LN), liver and lung. (**b**) Embedding of tumor cells from each tissue with ATs indicated by colored circles. (**c**) Representative histological sections of tumors and metastases taken for single-cell sequencing. (**d**) Stacked barplot showing percentage of tumor cells in each tissue committed to each archetype (colored) or uncommitted to an archetype (gray). (**e**) Stacked bar plot showing the cell cycle phase of cells committed to each archetype. (**f**) Enriched HALLMARK genesets associated with ATs from each tissue (indicated by colors). P-value is false discovery rate corrected, genesets where p > 0.1 are shown in gray. Enrichment represents log2 fold-change. P = primary, LN = lymph node, LI = liver, LU = lung.

Cellular heterogeneity in metastases followed a phenotypic continuum akin to that observed in primary tumors (Figure 4b, Supplementary Figure 4b-d). To characterize this continuum, AAnet defined five archetypes in the lymph node, four in the liver, and five in the lung, where all archetypes detected in the primary had an ortholog in at least two metastatic sites (Supplementary Figure 7b-c). Moreover, a highly conserved pattern of hallmark enrichment was observed in orthologous archetypes from different sites (Figure 4f, Supplementary Table 2). This demonstrates that the factors of variation contributing to cancer cell heterogeneity were largely recapitulated in primary tumors and metastases.

While orthologous archetypes were identified in many different metastases, analysis of archetypal affinity indicated their relative contribution to cellular heterogeneity in each tissue was somewhat distinct.

Lymph node metastases showed the strongest resemblance to primary tumors. All five archetypes identified in the lymph node had an ortholog in the primary tumor (Figure 4d-f). The proportions of cells committed to each archetype were also highly conserved, with the proliferative and immune-stimulatory archetypes the most abundant in both tissues (Figure 4d). Moreover, the changes in the relative abundance of the oxidative, hypoxic and cell death archetypes were marginal, collectively indicating cell heterogeneity is very similar between primary tumors and lymph node metastases.

Liver metastases differed from the primary and lymph node in the number and abundance of archetypes. Only four of the five archetypes identified in the primary tumor were detected in the liver, and more cells were committed to an archetype than in other tissues (Figure 4d-f). The absence of cells committed to the oxidative archetype was replaced by a greater proportion of cells committed to the proliferative and immune stimulatory archetypes, both nearly double their proportions in the primary tumor (32.94% and 23.54% respectively). Overall, while the archetypes were somewhat conserved with primary tumors and lymph node metastases, their relative contribution to cellular heterogeneity distinguished liver metastases from the other tissues.

Lung metastases showed the greatest difference from primary tumors and the other metastases. Five archetypes were detected in the lung, four of which were orthologous to archetypes in the primary tumor (Figure 4d-f). While still the most abundant, the proportion of cells committed to the proliferative archetype were significantly lower than in other tissues (18.36%). The oxidative archetype, enriched for OXPHOS and ROS hallmarks, was the second most abundant and almost equal to the proliferative archetype (17.72%). Conversely, the hypoxic archetype was not detected in the lung metastases. This is consistent with cancer cellular heterogeneity in these metastases being shaped by a oxygen-rich lung microenvironment.

The remaining archetype was unique to the lung. Markers of this archetype were significantly enriched for the cancer hallmarks of an oxidative metabolism (OXHPOS and ROS pathway) driven by genes encoding ETC components (ATP5F1A,B, NDUFA4,8,9, COX7C) and antioxidant enzymes (PRDX2,4,6) respectively (Supplementary Table 2). Additionally, hallmarks of TNF signaling via NF-κB, apoptosis, hypoxia, protein secretion, and coagulation were overrepresented, a profile of pathways indicating similarity to the immune stimulatory archtype. Importantly, enrichment of hypoxia was driven by a separate subset of glycolytic enzymes to those associated with the hypoxic archetype in other tissues, and did not include markers of HIF1A induction such as CITED2 (Supplementary Table 2).

Together, these analyses show that AAnet-defined archetypes in primary tumors are also identified in metastases. This suggests that factors influencing cellular heterogeneity are highly conserved between primary tumors and metastases. Differences between tissues were largely driven by the differential influence of these archetypes, potentially due to interactions with the metastatic microenvironment.

### Spatial Transcriptomics reveals organization and distinct cellular morphology of AAnet archetypes

To further validate the significance and biology of the archetypes identified by AAnet, we sought to investigate their structural organization within tumors. Thus, we performed spatial transcriptomics (Visium 10X) on tissue sections from two primary tumors that were grown *in vivo* for 8 weeks. These samples were collected from two of the tumors used for scRNAseq, with parts of these tumors frozen in OCT for sectioning. Gene expression was measured at 2,275 spatially distributed voxels in the two samples after QC (1,170 and 1,105 voxels respectively), with each voxel assaying expression in an area approximately 3-10 cells in size.

To map the archetypes from the scRNAseq to the spatial transcriptomic data, we first had to overcome the intrinsic differences in the data generated by these modalities. Raw data from these technologies is not directly comparable as they have very different sample processing protocols (digestion and droplets vs freezing and sectioning) and biological resolutions (single cells vs multiple cells). These differences create batch effect, where noise introduced in the process of data generation dominates the biological differences between samples. This batch effect was evident when spatial voxels were embedded with the scRNAseq data from the primary tumor, as there was little alignment between the modalities despite being generated from matched biological samples (Supplementary Figure 8a).

To address this, we leveraged our single-cell multi-modal generative adversarial network (scMMGAN) [4] (Methods). This approach uses adversarial learning to align data between modalities, enabling integrated downstream analysis while preserving data geometry (Figure 5a). We used scMMGAN to generate a scRNAseq measurement for each spatial voxel (scVoxels) that is aligned to our primary scRNAseq dataset. We can then input the aligned spatial data into the AAnet encoder trained on the primary scRNAseq data. This provides an archetypal representation for each spatial voxel based on our previously characterized primary archetypes (Figure 5c, Supplementary Figure 8b-c). The ability to extend archetypal analysis to previously unseen data through the neural network framework is an important feature of AAnet versus existing archetypal analysis approaches.

**Figure 5.**
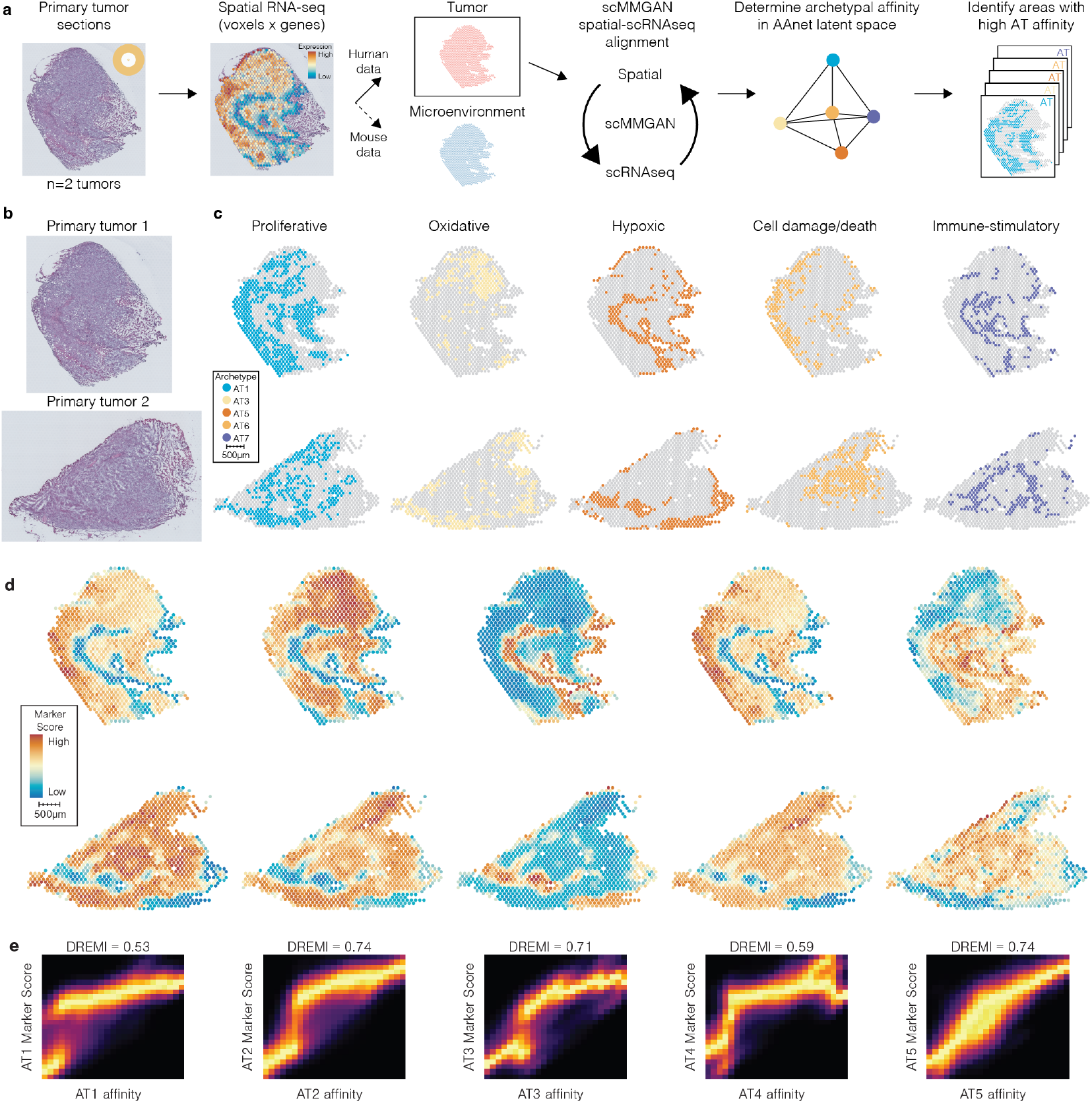
(**a**) Approach used to define a spatial map of TNBC tumor archetypes using scMMGAN and AAnet. (**b**) Histological sections of TNBC xenografts used for spatial transcriptome sequencing (n=2). (**c**) Spatial voxels with a strong affinity (>0.3) for each archetype (colored) as determined by scMMGAN and AAnet. (**d**) Spatial plots colored by marker geneset score for archetype. (**e**) Density resampled estimate of Mutual Information (DREMI) score between archetypal affinity and marker geneset scores.

With spatial transcriptomic data embedded as scVoxels in the AAnet latent space, we calculated their affinity to each archetype which was mapped back to their corresponding spatial location in the primary tumor (Methods, Supplementary Figure 8). Archetypal affinities and marker geneset expression scores shared high mutual information across voxels (Methods, Figure 5d-e). This indicates that biology that defined each archetype had been retained in the spatial mapping process.

Archetypes showed spatial organization and were associated with distinct cell morphologies in primary tumors:

### Proliferative archetype (blue)

Voxels with high affinity for the proliferative archetype (affinity > 0.3) formed the bulk of the tumor yet were markedly absent from the central areas of each tumor section. Areas with high affinity for this archetype were enriched for cycling cells (Supplementary Figure 5c). These findings were consistent with the scRNAseq commitment analysis (Figure 3e) indicating that proliferation was a dominant factor of cell heterogeneity in the primary samples.

### Oxidative/adipogenic archetype (yellow)

Areas of the tumor with high affinity for the oxidative archetype were located in close proximity to the proliferative archetype (Figure 5d). The affinity scores and expression of marker genesets were also significantly correlated (Figure 5e-f), indicating a relationship between oxidative metabolism and a proliferative cell state.

### Hypoxic archetype (orange)

Strikingly, areas with a strong affinity for the hypoxic archetype were localized to central and peripheral regions of primary tumors, devoid of the proliferative and oxidative archetypes (Figure 5c). These areas showed enriched expression of markers associated with oxygen-independent glycolysis and ribosomal subunits (Figure 5d-e).

### Cell damage/death archetype (amber)

The cell death archetype localized to areas with high expression of mitochondrial genes and was strongly correlated with the proliferative and oxidative archetypes (Figure 5c-e). While preferential enrichment of mitochondrially encoded genes is often used as a marker of cell death, mitochondrial genes encode critical components of the electron transport chain necessary for oxidative energy production. Consequently, the association between this archetype and the proliferative/oxidative archetypes may identify oxygen-rich areas within primary tumors rather than cell death.

### Immune-stimulatory archetype (purple)

Notably, affinity for the immune-stimulatory archetype was highest surrounding the hypoxic archetype within the tumor. These areas showed enriched expression (Figure 5c) of cytokines and antigen presenting proteins, such as CXCL1 and HLA-A/B/C/B2M (Figure 5d-e), consistent with the hallmark associations of this archetype. Interestingly, the immune-stimulatory archetype appears to demarcate the hypoxic and proliferative cancer cell archetypes.

Together, this analysis shows identified archetypes have unique and distinct spatial organization within the tumor, further validating the ability of AAnet to identify unique cellular biology and structure of cancer cells within a phenotypic continuum of cell states. Further, the organization of the AAnet archetypes is consistent with a model whereby the local microenvironment may play a critical role in determining the organization of cellular heterogeneity in primary tumors.

### Archetypes are associated with distinct cell types and metabolic niches in the microenvironment

With clear spatial organization of cancer cell archetypes derived by AAnet within the tumor, we next sought to determine if, beyond spatial location, the tumor microenvironment may also be playing a role in archetypal development and commitment. We therefore assessed if microenvironmental cells were spatially structured into meaningful microenvironmental archetypes (ME-ATs) (Figure 6a). The xenograft model enabled separation of expression at each voxel into tumor and ME data based on alignment to the human or mouse genome, respectively. Then, we used AAnet to deconvolute the murine data into ME-ATs to investigate archetype-specific cell types and biological processes.

**Figure 6.**
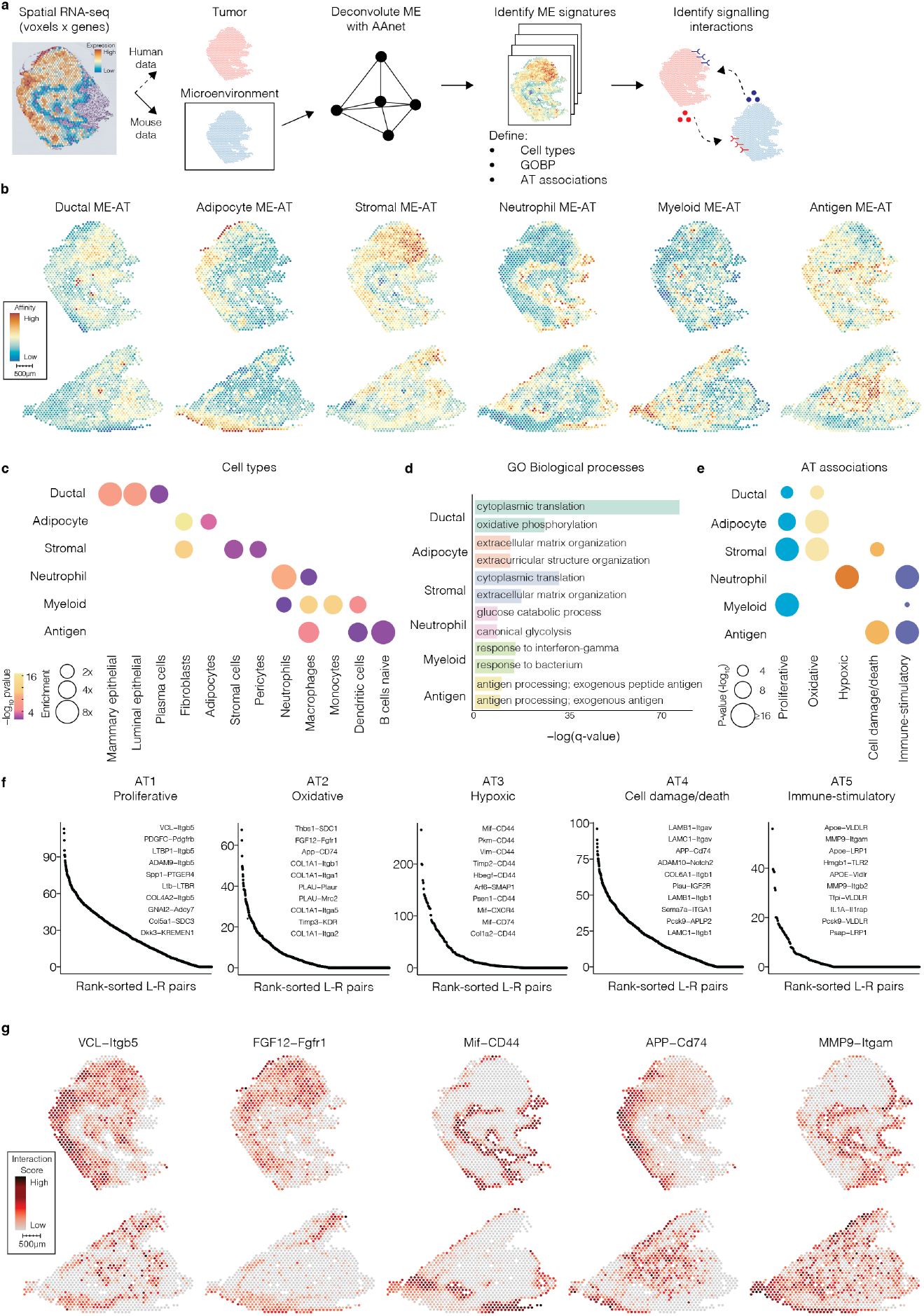
(**a**) Approach to defining tumor microenvironments (ME) associated with key archetypes (AT) using AAnet. (**b**) Microenvironment-archetype (ME-AT) affinity scores for each spatial voxel. (**c**) Enrichment of cell-types associated with each ME-AT. P-value is false discovery rate corrected. Enrichment represents log2 fold-change. (**d**) Enrichment of top 2 biological processes from the gene ontology (GO) associated with each ME-AT. Processes are ranked by FDR (-log10). (**e**) ME-ATs (y-axis) significantly associated with each tumor AT (x-axis). Color reflects tumor AT, enrichment represents log2 fold-change. (**f**) Top ranked ligand-receptor pairs expressed in spatial voxels with a strong affinity for each archetype and their colocalized microenvironments. Capitalized gene symbols indicate genes with expression in tumor cells (human). Title case symbols indicate genes with expression in ME cells (mouse). Pairs are ranked by FDR (-log10). (**g**) Ligand-receptor interaction score across spatial voxels for top pairs associated with each archetype.

AAnet defined six ME-ATs with unique patterns of spatial organization (Figure 6b, Methods). Each ME-AT is enriched for specific cell types and biological processes (Figure 6c-d). These include a ductal ME-AT with high enrichment of mammary epithelial cell markers in voxels overlaying to breast ducts, as well as an adipocyte ME-AT enriched for adipocyte markers and most highly expressed on the margins of the tumor sections with residual mammary fat pad. This correspondence with underlying histological features provides orthogonal validation for AAnet. AAnet also identified a stromal ME-AT, enriched for fibroblasts, capillary-lining pericytes, and stromal cells, as well as biological processes related to ECM organization, oxidative phosphorylation and angiogenesis; a neutrophil ME-AT, enriched for neutrophils and biological processes related to glycolysis, leukocyte chemotaxis and hypoxia; a myeloid ME-AT enriched for markers of monocytes, macrophages and dendritic cells, with an enriched response to interferon-gamma, immune effector processes and leukocyte mediated cytotoxicity; and an antigen-response ME-AT enriched for macrophages and genes involved in antigen processing and presentation of exogenous peptide antigen via MHC class II, and immunoglobulin-mediated immune response (Figure 6c-d). Therefore, AAnet deconvolutes expression in the microenvironment into stromal and immune components.

Next, we explored the spatial relationship of the ME-ATs to the cancer cell ATs and uncovered specific associations (Figure 6e, Methods). Areas of the tumor with a high affinity for the proliferative AT were strongly associated with the myeloid and stromal ME-ATs, while the oxidative/adipogenic AT showed preferential enrichment for the stromal ME-AT. This suggests these archetypes colocalize with highly metabolic and vascularized microenvironments. Both of these archetypes were also associated with the ductal and fat-pad ME-ATs, concordant with the localization of these normal mammary gland structures in the histological sections. Notably, the hypoxic AT colocalized specifically with the neutrophil ME-AT. These interactions occurred at internal parts of the tumor section, indicating that these archetypes are associated with decreased oxygen availability and increased neutrophil chemotaxis. The cell death AT was strongly associated with the antigen ME-AT, indicating an active presentation of cancer cells to the immune system, cancer cell death and phagocytosis of dead and dying cells by macrophages. Interestingly, the immune-stimulatory cancer cell AT, defined by its enrichment for gene sets related to immune signaling, was associated with the antigen and neutrophil ME-ATs, indicating a strong overlap in signaling and cell phenotype between the cancer and microenvironmental cells. Thus, we observed multiple examples of spatially-localized phenotypic mimicry between cancer and microenvironmental cells, most notably in metabolic and immune signaling.

To investigate direct cell-cell signaling mechanisms for phenotypic mimicry, we analyzed ligand-receptor pairs (LR-pairs) for evidence of paracrine interactions (Methods). We again used our hybrid model system to delineate between tumor (human) and ME (mouse) expression, allowing us to establish the direction of signaling. Specifically, LR-pairs with a ligand expressed in the human data and its cognate receptor expressed in the mouse data indicate signaling from the tumor to the microenvironment, and vice versa (Figure 6f). We first determined the coexpression of annotated LR-pairs across all voxels and calculated their enrichment in areas with a high affinity for each archetype. For all archetypes, the proportion of ligands originating from the tumor and the microenvironment were approximately 50%. However, the strongest LR-pairs that localized to high affinity regions differed between archetypes.

Tumor cells expressed stromal growth factors in areas with a high affinity for proliferative and oxidative/adipogenic archetypes. These included platelet-derived growth factor (AT Ligand: PDGFC, ME Receptor: Pdgfrb) and fibroblast growth factor (AT Ligand: FGF12, ME Receptor: Fgfr1) pathways that promote blood vessel formation [12]. The hypoxic archetype was characterized by interactions between CD44 expressed in tumor cells and a variety of ligands expressed in the microenvironment, including Mif, which may enhance neutrophil accumulation within the hypoxic AT. Examples of cancer-microenvironmental cross talk in the cell death AT are evidenced by App-Cd74, where Cd74 is an MHCII molecule, and sema7a-ITGA1, where sema7a is a potent immune modulator. We also observe strong overlap in LR-pairs regulating immune-cancer cell crosstalk in the immune-stimulatory AT including MMP-itgam, IL1A-Ilr1rap, and Hmgb1-TLR2. (Figure 6g).

Together, these results highlight the spatial colocalization of distinct cancer archetypes with unique microenvironments, where paracrine interactions that may enhance phenotypic mimicry are an important determinant of intratumoral heterogeneity.

### Intratumoral metabolic heterogeneity in TNBC and alignment to human breast cancers Figure 7

The discrete localization of distinct metabolic phenotypes within the tumor, including the oxidative phosphorylation-enriched proliferative AT versus the glycolytic-enriched hypoxic AT (Figure 3d and Figure 4f), as well as the metabolic mimicry of the microenvironmental cells in those regions (Figure 6d), led us to ask if targeting a specific metabolic program might impact tumor growth. To delve into this question, we first examined the metabolic state of the microenvironments colocalized with each archetype by comparing genes associated with glycolysis and the TCA cycle in each cancer AT and associated microenvironment. Indeed, the concordance between the cancer AT and the microenvironment was driven by the correlation in expression of metabolic enzymes (Figure 7a). Enzymes in the tricarboxylic acid cycle (TCA-cycle), a pathway which requires oxygen to generate energy, were uniformly highly expressed in the tumor and ME of the proliferative, oxidative and cell death archetypes. In contrast, glycolytic enzymes were clearly enriched in the hypoxic AT and associated ME cells (Figure 7a). Enzymes most highly enriched in the hypoxic area were associated with oxygen-independent glycolysis in tumors [1] and include PDK1, an inhibitor for the entry of pyruvate into the TCA cycle. This indicates that the metabolic heterogeneity in primary tumor archetypes is mirrored in their local microenvironments, and the intratumoral hypoxic regions of the tumor are driven by glycolysis ending in the accumulation of lactate. Interestingly, hypoxic niches are known to provide a permissive environment for maintenance of both pluripotent stem cell and cancer stem cell populations [34, 53]. AAnet identified that SLC2A1 (GLUT1) and SLC2A3 (GLUT3) are enriched in cancer cells that reside in that niche.

**Figure 7.**
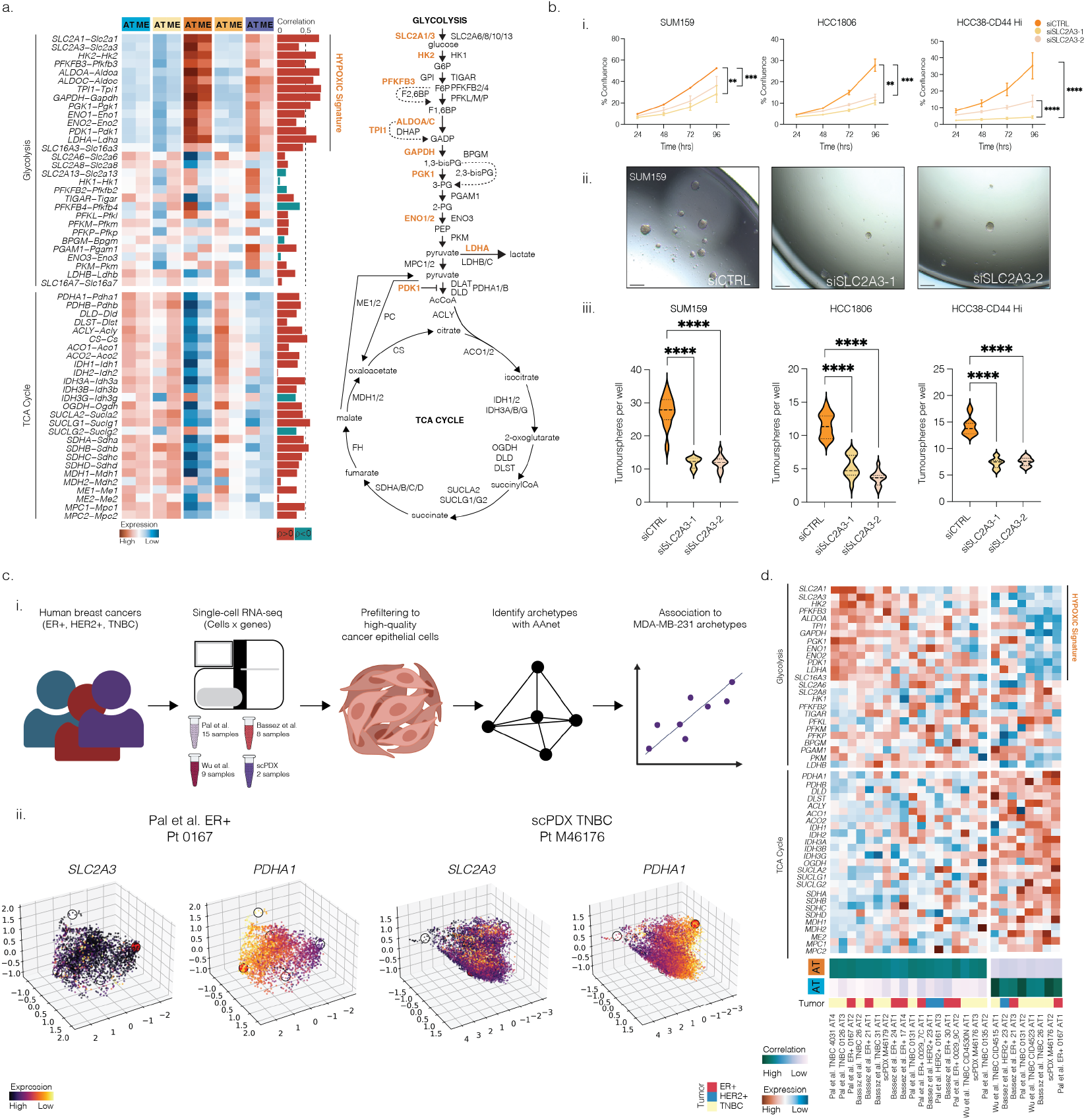
**(a)** Heatmap showing expression of metabolic genes (glycolysis and oxidative phosphorylation) across cancer archetypes and associated microenvironmental cells. **(b)** i. Cell growth of HCC1806, SUM159 and HCC38-CD44 Hi cells treated with either control siRNA (siCTRL) or siRNA targeting SLC2A3 (siSLC2A3-1, siSLC2A3-2) over a 96-hour period. *n =* 3 independent experiments, triplicate wells analyzed per condition. Statistical significance defined by one-way ANOVA. ii. Representative images of SUM159 tumorspheres treated with control or SLC2A3 targeted siRNA. iii. Quantitation of tumorsphere-forming capacity in control (siCTRL) and SLC2A3 knockdown cells (siSLC2A3-1 or siSLC2A3-2) in SUM159, HCC1806 an HCC38-CD44 Hi cells. *n* =3 independent experiments, 12 wells analyzed per condition, statistical significance defined by ordinary one-way ANOVA with Tukey’s multiple comparison post-hoc analysis. **(c)** i. Diagrammatic representation of human breast cancer sample single-cell analysis. ii. Visualization of cancer epithelial cells from two patients, colored by markers for oxygen-independent glycolysis (*SLC2A3*) and entry into the TCA cycle (*PDHA1*). **(d)** Heatmap showing expression of metabolic genes from (a) across 26 human cancer archetypes, 18 (left) associated with the hypoxic archetype and 8 (right) associated with the proliferative archetype. Archetypes are from breast cancer samples across three different breast cancer subtypes.

Given that GLUT1 is ubiquitously expressed in normal cells throughout the body, we asked if ablating GLUT3, whose expression is largely confined to the brain and sperm in normal tissues, could eradicate the aggressive phenotype of the cancer cells in the hypoxic niche. In addition, GLUT3 expression is increased in TNBC and associated with metastasis and poor prognosis [44]. Three TNBC cancer stem cell-enriched cell lines (SUM159, HCC1806 and HCC38-CD44Hi) were treated with a control siRNA (siCTRL) or two independent siRNAs targeting SLC2A3/GLUT3 (siSLC2A3-1 and siSLC2A3-2). Efficient knockdown of SLC2A3 was confirmed by qPCR (Supplementary Figure 9). SLC2A3 knockdown significantly inhibited cell proliferation in all cell lines tested (Figure 7b, Supplementary Figure 9). Excitingly, we confirm that SLC2A3 knockdown significantly inhibits tumorsphere formation, an *in vitro* surrogate assay for *in vivo* tumor-initiating ability (Figure 7b). Together, these results suggest that SLC2A3 is critical for maintenance of the cancer stem cell phenotype in TNBC and add to previous data indicating a role for GLUT3 in EMT and migration [44].

To examine the clinical relevance of the identified archetypes, we analyzed single-cell transcriptomes from human breast cancer cells across four distinct studies, corresponding to 34 samples from three major breast cancer subtypes (ER+, HER2+, TNBC) (Figure 7c) [6, 35, 51]. First, AAnet identifies 155 archetypes across the 34 samples and shows interesting similarities and differences across samples, cohorts, and breast cancer subtypes (Methods, Supplementary Figure 10). To further investigate the association of human archetypes with the proliferative AT (blue) and hypoxic AT (orange), we identify 18 archetypes across all human tumors that have transcriptomic profiles similar to AT3 (cosine similarity > 0.25) and dissimilar to AT1 (cosine similarity < −0.25), as well as 8 archetypes similar to AT1 and dissimilar to AT3 (Figure 7d). These metabolic archetypes are common in breast cancer, represented in 26 archetypes spanning 20 human tumors. Similar to the xenograft model, 6 tumors contain both a hypoxic AT and a proliferative AT within the same sample. Together, these results show correspondence between the metabolic profiles of archetypes identified by our model and human breast cancer tumors, and further, the identified archetypes are relevant across breast cancer subtypes.

Visualization of the metabolic markers from Figure 7a reveals, for human archetypes associated with hypoxia, enrichment for the hypoxic signature within the glycolysis pathway and low expression of genes related to the TCA cycle. Conversely, human archetypes similar to the proliferative archetype showed low expression for hypoxic genes and higher enrichment for TCA cycle genes.

Notably, there is no significant enrichment for a particular cancer subtype and association with AT1 or AT3 (KS test p>0.05 for all tests), nor significant difference between cancer subtypes in proportion of cells committed to hypoxic or proliferative archetypes (Wilcoxon rank sums test p>0.05 for all tests). These data show that AAnet can be used to identify phenotypic similarities across cancer subtypes, and thereby offers a functional method beyond hormone and molecular subtyping to classify breast cancers for therapeutic targeting.

## Discussion

It is now recognized that non-genetic programs (e.g., epigenetic, transcriptional, translational) are a major driver of tumor heterogeneity. The dynamic and reversible nature of non-genetic heterogeneity likely favors rapid evolution of cancer cell states (e.g., seconds, minutes or hours) to enable survival in unfavorable microenvironments encountered throughout the metastatic cascade and in response to therapy. Thus, as single-cell technologies continue to resolve the breadth and structure of non-genetic heterogeneity in cancer and stromal cells within and across patient tumors, developing strategies to identify and validate the specific cell states and molecular mechanisms that fuel cancer progression remains a significant technological and biological challenge.

To address this knowledge gap, we developed AAnet, an archetypal analysis method to identify archetypal cell states within and between samples and their associated biological processes. Archetypal analysis is a framework to describe a dataset as a convex combination of extreme, or archetypal, observations. In contrast to other unsupervised approaches to characterize such data, archetypal analysis is aptly suited for both identifying key cell states reflecting distinct biological processes and analyzing the cellular state space as a continuum of cells committed to these processes. However, identifying the archetypal states remains a fundamental challenge of archetypal analysis. In particular, nonlinearities can worsen the performance of existing archetypal analysis tools, as the extreme states of the data geometry do not conform to the extreme states of the data space.

AAnet solves this problem by learning a transformation of the data into a simplex, rather than fitting a simplex on the data directly. The latent space of the autoencoder thus preserves relationships between cells and characterizes cells by their commitment to each archetype. We further regularize the latent space to initialize the archetypes to diffusion extrema (inferred from the cell-cell affinity graph) for improved accuracy and robustness. Applied here to single-cell data from pre-clinical and clinical breast cancer samples, we have shown that AAnet enabled the discovery of biologically and functionally-distinct archetypes within a phenotypic continumm of cell states within the tumor and captured their associated molecular drivers. First, in a pre-clinical xenograft model comprising primary tumors matched with lung, liver, and lymph node metastases, we identify six unique archetypes across primary and metastatic tissues, with each archetype defining unique biology of cells committed to that extrema. Interestingly, we show that the number and distribution of cells committed to archetypes in primary tumors is remarkably similar to those found in lymph node metastases, yet liver metastases differ by the loss of one archetype, and the lung metastases deviate from primary, lymph node and liver metastases via the emergence of one new archetype. These analyses demonstrate that AAnet can reveal the emergence of new cell states, the number of cells committed to a specific archetype, and the underlying biology that facilitate site-specific metastatic adaptation.

Critically, we validate the significance of the archetypes identified by mapping the scRNAseq data to matched spatial transcriptomic data via scMMGAN [4]. These data confirm that AAnet-defined archetypes resolve into distinct spatially-localized regions within the tumor. Further, we show that AAnet has revealed a unique perspective on the organization of the associated microenvironments. Specifically, each archetype is enriched with distinct stromal cell types; for example, the proliferative archetype is enriched with fibroblasts, hypoxic archetype with neutrophils, and immune-stimulatory archetype with macrophages and dendritic cells. Thus, AAnet robustly identifies functional and spatially distinct cellular archetypes within a tumor.

Of note, we uncovered metabolic heterogeneity not seen before in TNBC, where cells in discrete archetypes utilize distinct metabolic programs. We have recently analyzed bulk RNA-seq data (METABRIC and TCGA) and shown that TNBC exhibit a unique highly metabolic gene expression phenotype, upregulating a range of pathways,including glycolysis, compared to other breast cancer subtypes such as Luminal A [37]. Our archetypal analysis provides further granularity to these data, showing the individual contributions of each archetype to this unique TNBC metabolic signature. For example, the hypoxic archetype clearly contributes to the high expression of SLC2A1, SLC2A3 (GLUT3), HK1, HK2, ALDOA, ALDOC, TPI, GAPDH, PGK1, ENO1, PDK1 and LDHA in the bulk RNAseq data, with contributions also from the immune-stimulatory archetype but not from the most abundant proliferative archetype. By comparison, these findings reveal that different regions of the tumor uniquely invoke glycolysis or oxidative phosphorylation, which could not be determined when analyzing existing bulk RNA-seq data. Furthermore, we show that we could use our identified archetypes to predict novel therapeutic targets within distinct cellular subsets, such as the glucose transporter GLUT3 in stem cells within the hypoxic archetype. Interestingly, we also discovered that the distinct metabolic phenotypes of two archetypes are strikingly reflected in the microenvironmental cells associated with those archetypes. Importantly, we also found these distinct archetypes are present in human breast cancer samples. GLUT3/SLC2A3 expression was present at highest levels in the hypoxic archetype, with high levels also seen in the surrounding immune archetype. A previous study has suggested a role for GLUT3 in regulating the inflammatory microenvironment in TNBC [44], which may also play a role in the matched metabolic phenotypes of the tumor and surrounding microenvironment cells.

Tumor heterogeneity remains a significant clinical challenge for diagnosis and therapeutic management. The discovery of the AAnet archetypes in scRNAseq data has enabled segmentation of tumors into functionally distinct regions comprising cancer cell states associated with unique cellular microenvironments. Together, these data resolve tumor heterogeneity to a level not yet achieved with previous computational tools. In the future, classifying patients according to biological archetypes with tools like AAnet is likely to improve tumor sub-classifications. Applied to samples before and after specific treatment, we can begin to learn how archetypes change over time, in response to specific therapies, and in different metastatic sites. Moreover, this approach will reveal the molecular programs driving each cellular archetype, as well as when and how they emerge. Ultimately, these tools will deliver the knowledge to enable the development of improved and effective therapeutic strategies.

Importantly, AAnet is a flexible framework that can be used both independently and as a part of a large single-cell analysis pipeline to interpret the archetypal distribution underlying any single-cell dataset. Given its widespread utility and generalizability to characterize cells, AAnet is a valuable tool for the single-cell community.

## Supplemental Figures

**Supplementary Figure 1.**
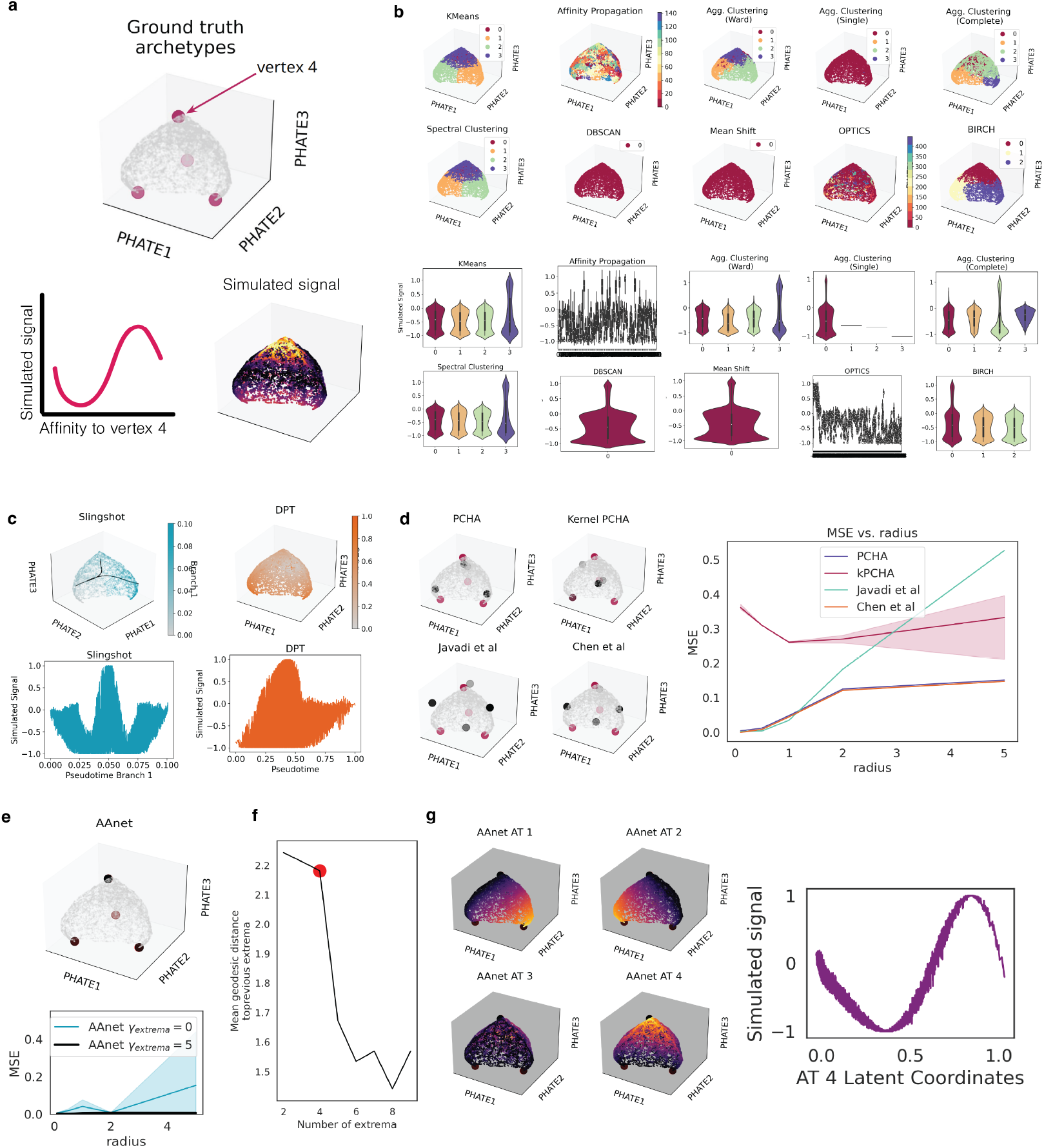
Curved Tetrahedron example. (**a**) Simulated curved tetrahedron, colored by signal defined with respect to affinity to archetype 4 (vertex at top). (**b**) Cluster assignments and signal comparison for 10 clustering algorithms. (**c**) Trajectories and signal comparison from Slingshot with KMeans clusters as input (left) and diffusion pseudotime (right). (**d**) Inferred archetypes from each archetypal analysis method (black) versus ground truth archetypes (red) and mean squared error between real and ground truth archetypes over increasing curvature. (**e**) AAnet-learned archetypes and mean squared error between real and ground truth archetypes over increasing curvature. (**f**) AAnet-inferred number of archetypes. (**g**) AAnet-learned archetypal coordinates. AAnet recapitulates sine signal with respect to archetype 4’s latent coordinates.

**Supplementary Figure 2.**
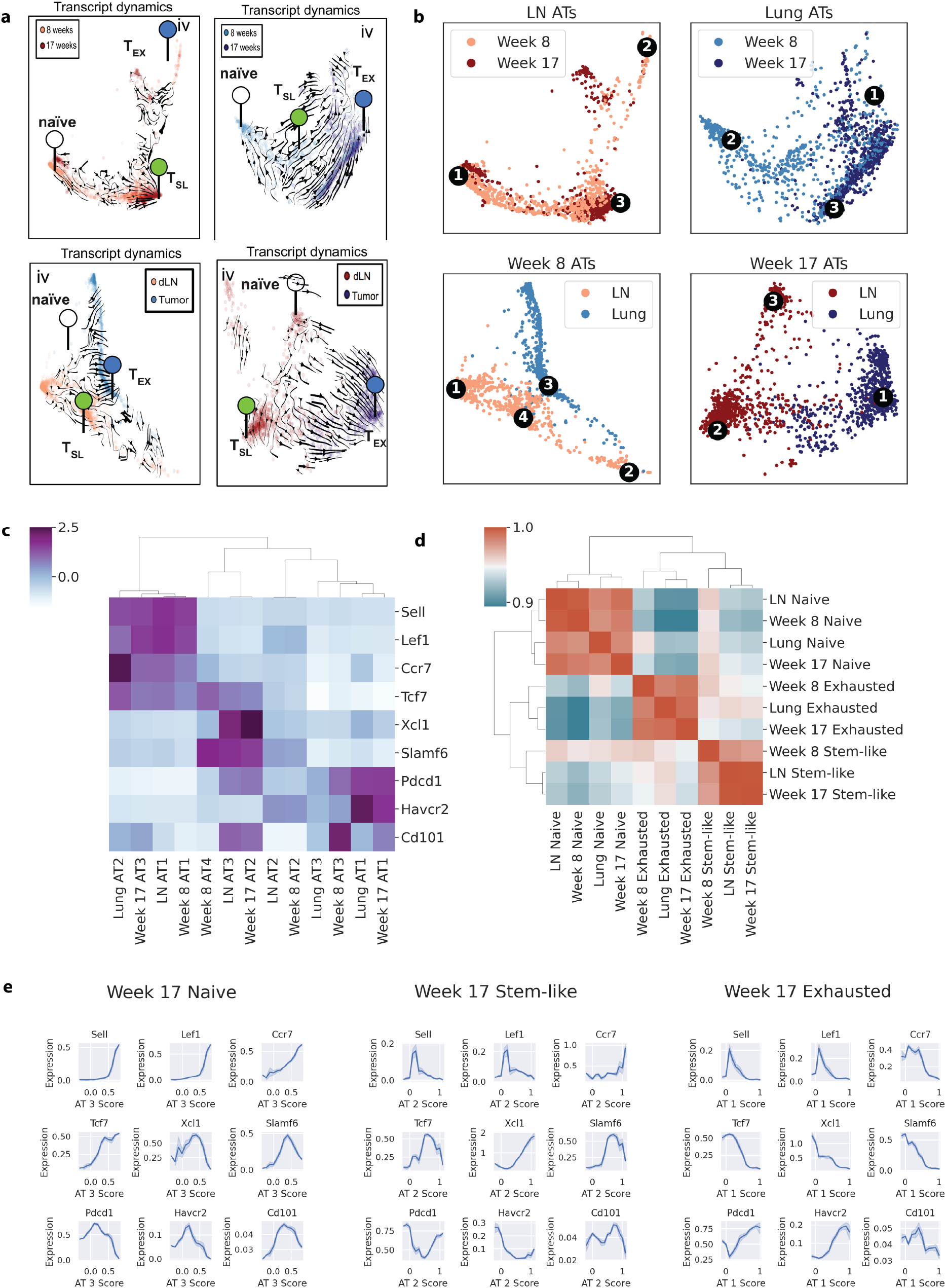
AAnet on CD8+ T cells. (**a**) Hand-annotated archetypes from [10]. (**b**) AAnet-learned archetypes for each embedding. (**c**) Normalized expression for each archetype for key annotation genes. (**d**) Cosine similarity between archetypes for all measured genes. (**e**) Expression over archetypal coordinates for Week 17 embedding.

**Supplementary Figure 3.**
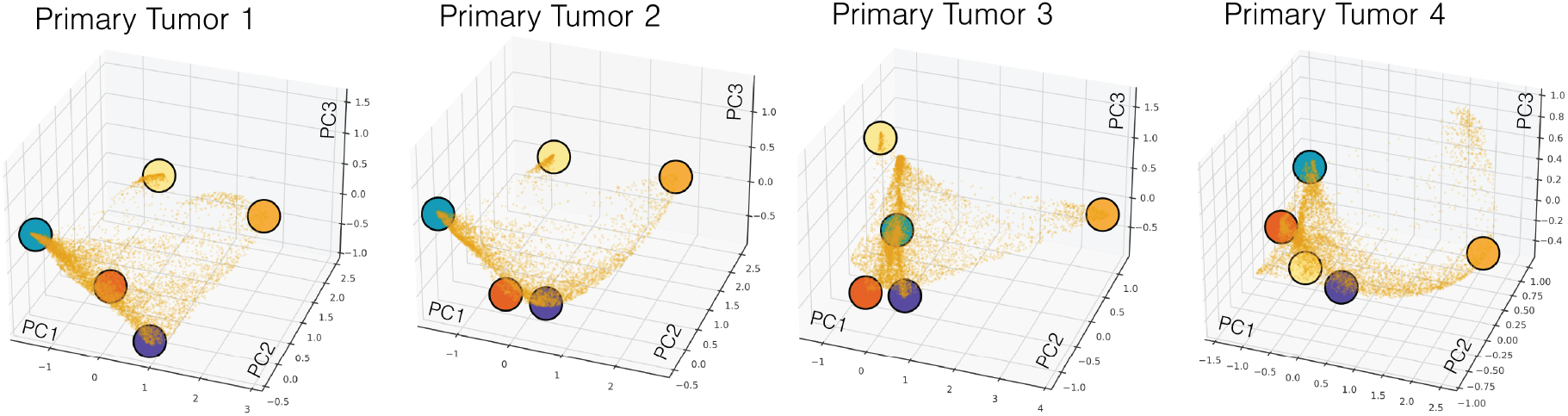
Primary tumor replicates independently characterized with AAnet. Archetypes colored based on orthologous archetype in combined embedding (Figure 3).

**Supplementary Figure 4.**
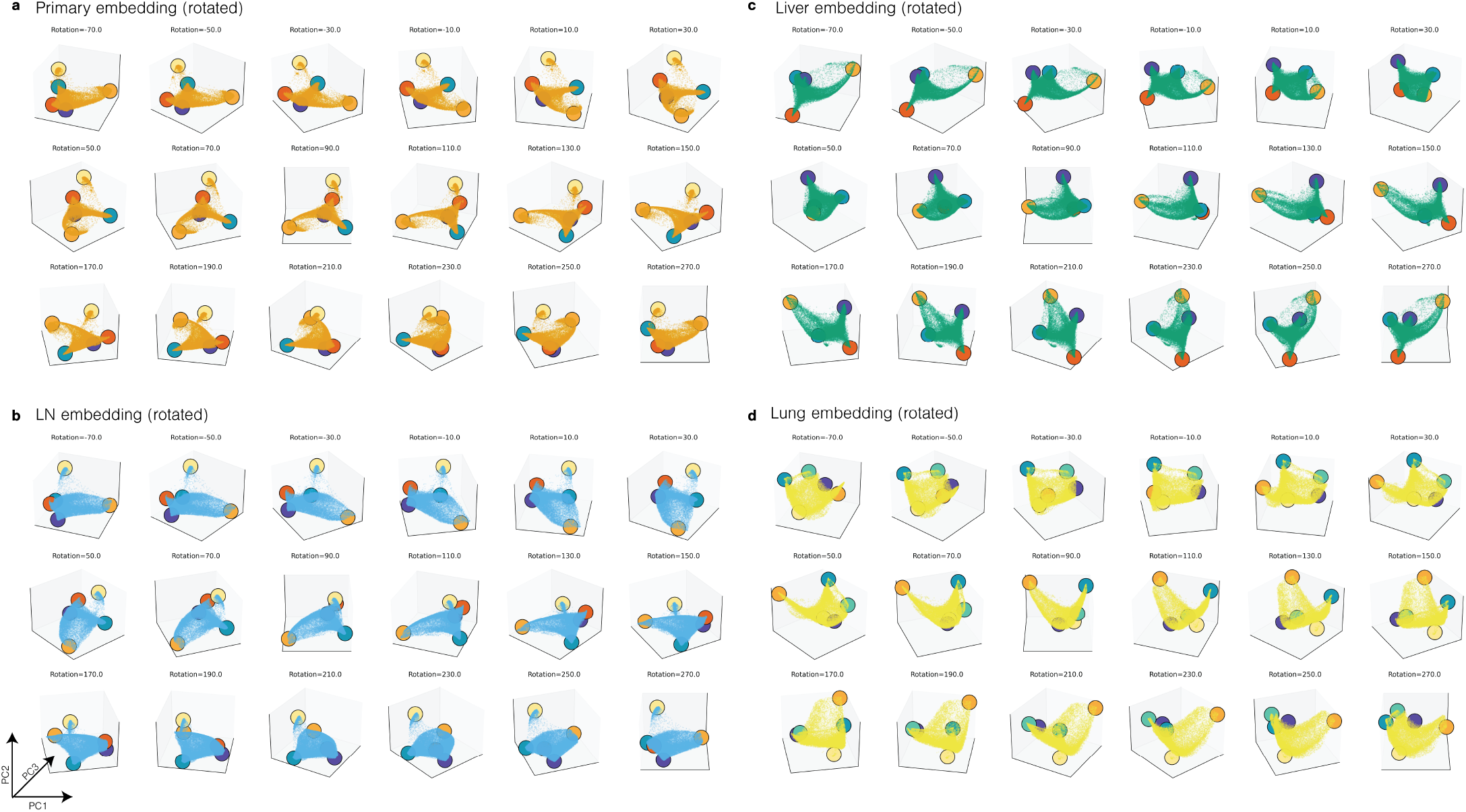
Cell embedding rotation. PCA embedding rotated around PC3 for (**a**) Primary (**b**) LN (**c**) Liver (**d**) Lung.

**Supplementary Figure 5.**
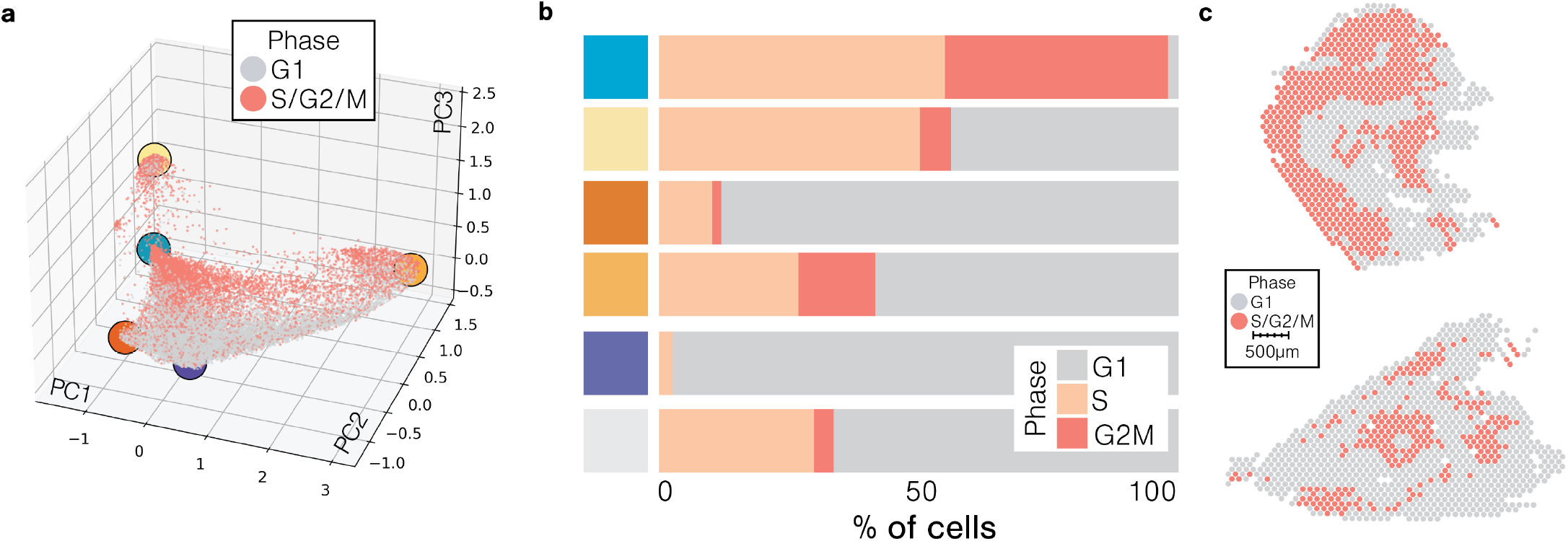
Cell cycle characterization. (**a**) Cell cycle commitment based on competitive gene set enrichment from scRNAseq data. (**b**) Cell cycle commitment of cells closest to each archetype. (**c**) Commitment of each spatial voxel to each cell cycle phase.

**Supplementary Figure 6.**
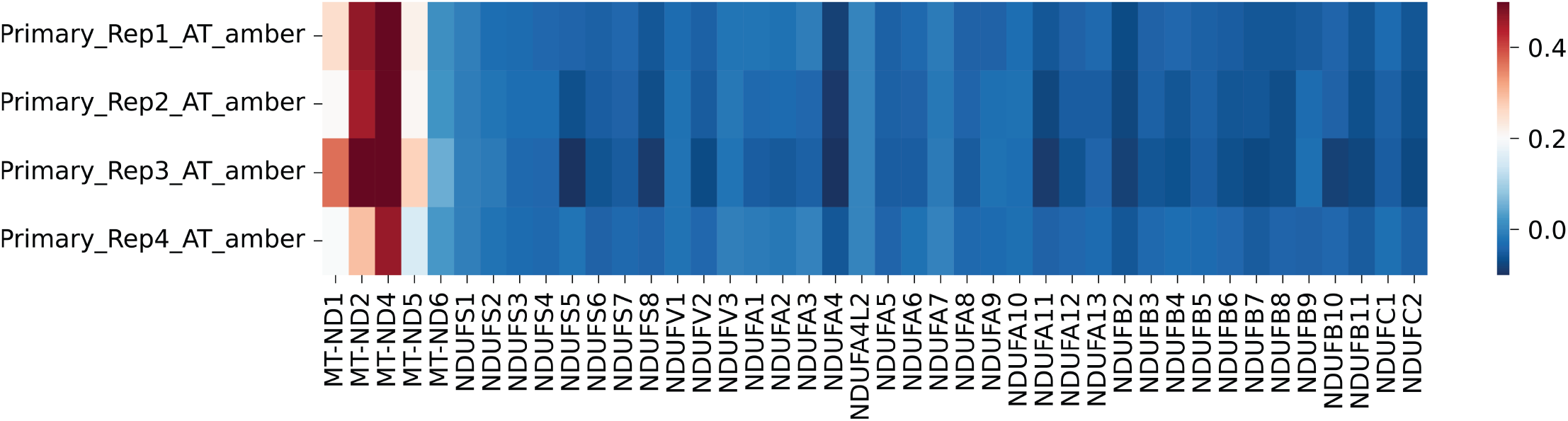
Mitochondrial expression. Expression of mitochondrially and somatically-encoded electron transport chain genes in cell damage/death (amber) archetypes from primary replicates.

**Supplementary Figure 7.**
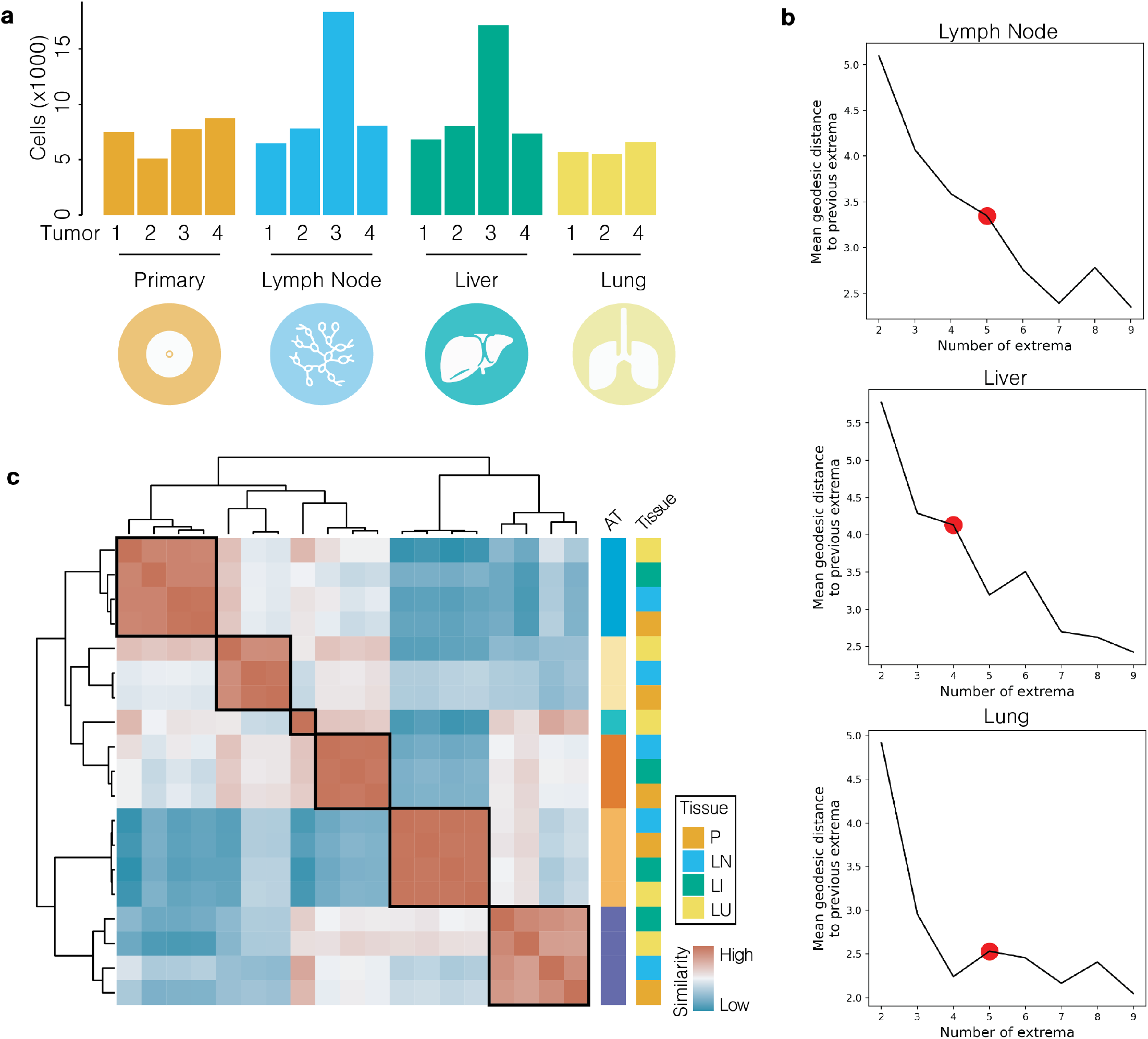
Comparison with metastatic tumor samples. (**a**) Number of cells from each tumor for each tissue. (**b**) Number of archetypes for each tissue. (**c**) Cosine similarity characterizing relationships between archetypes.

**Supplementary Figure 8.**
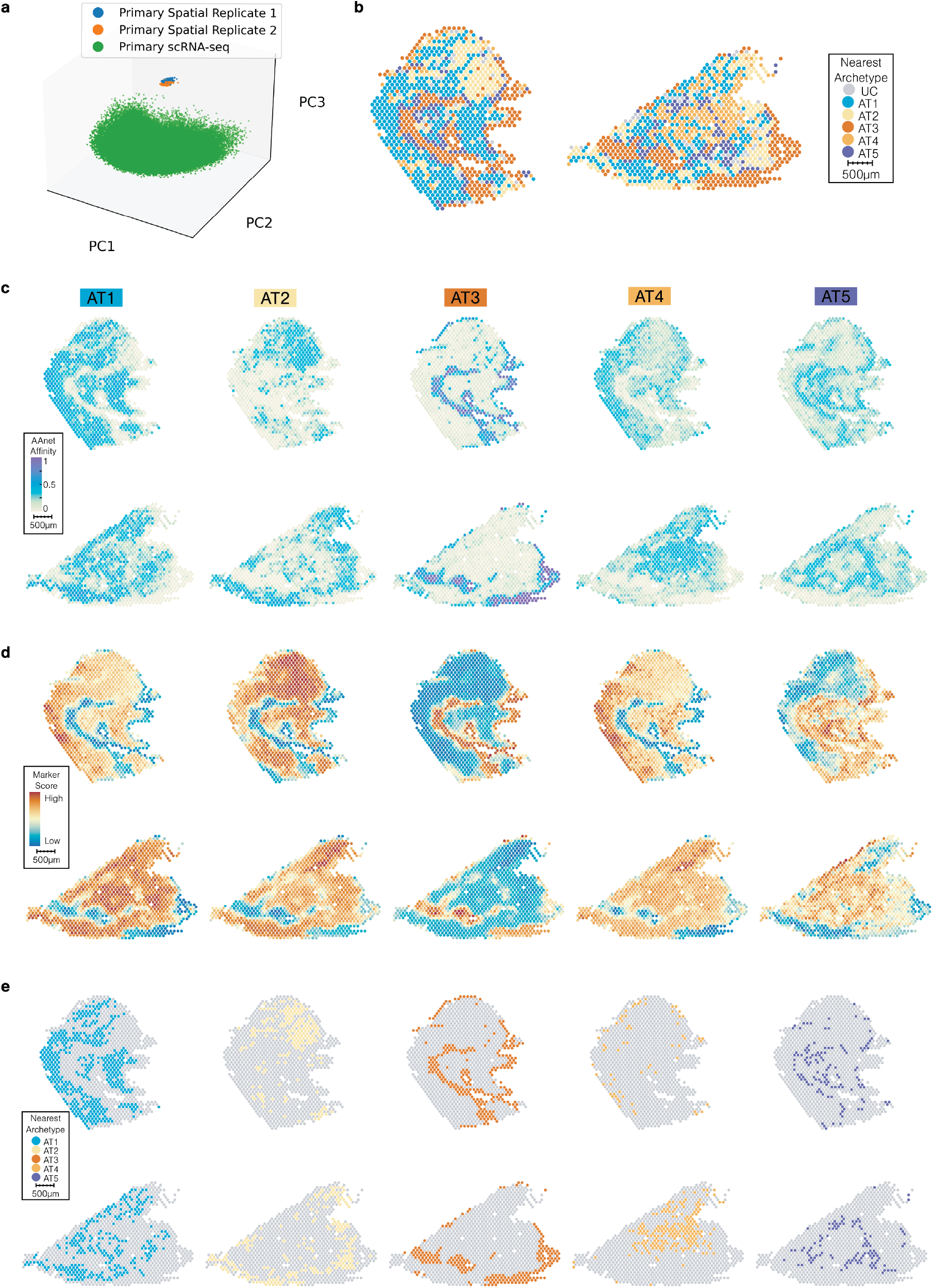
Spatial archetypal and gene set characterization. (**a**) Embedding without scMMGAN alignment shows batch effect. (**b**) Overall commitment, where voxels that remained uncommitted colored gray. (**c**) Archetypal affinity for each archetype after scMMGAN alignment. (**d**) Core gene set enrichment for each archetype. (**e**) Commitment of each spatial voxel to each archetype.

**Supplementary Figure 9.**
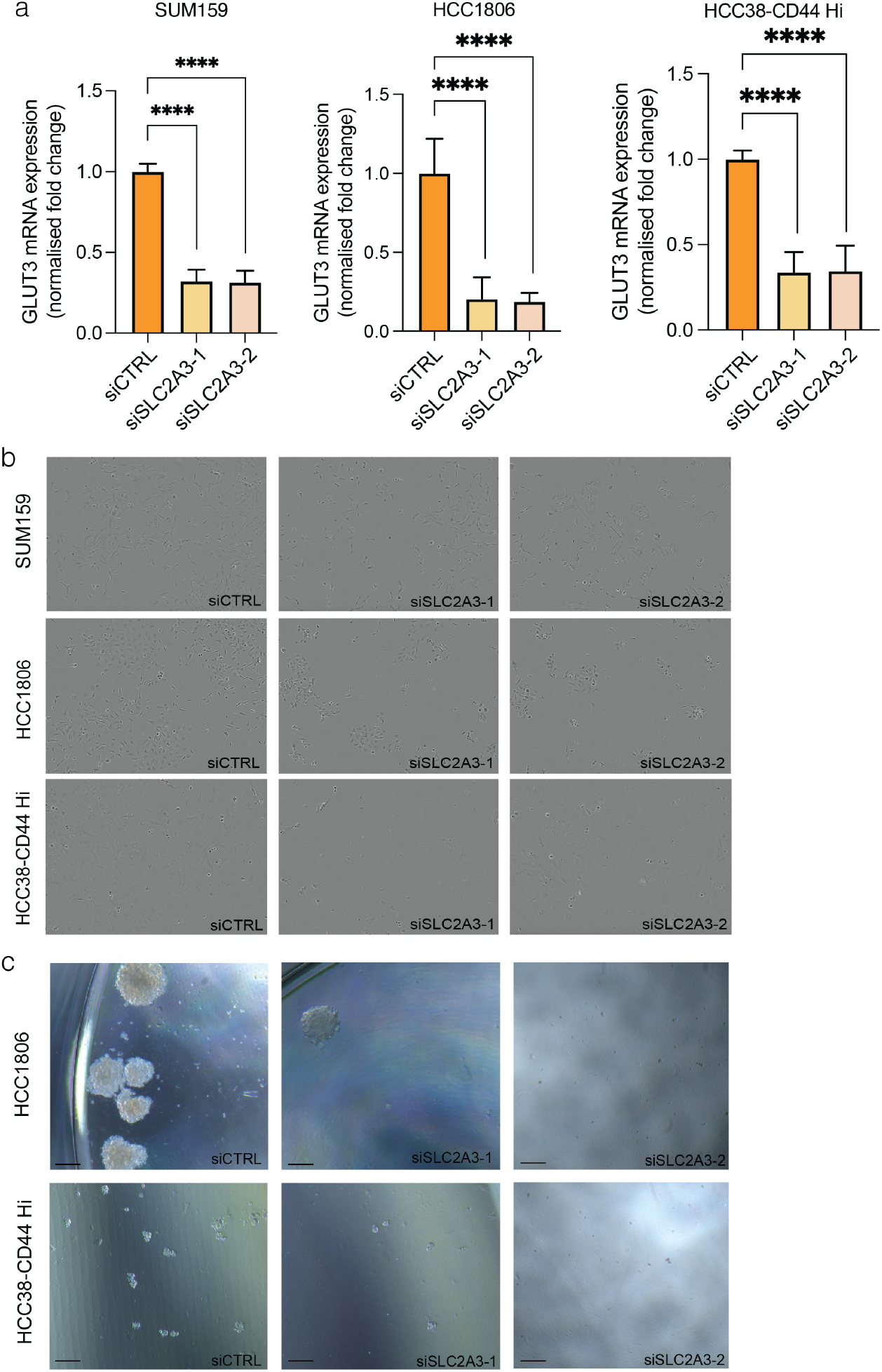
Metabolic heterogeneity and alignment to human breast cancers. (**a**) Quantitative PCR analysis of SLC2A3 mRNA expression following control or SLC2A3 targeted siRNA treatment in SUM159, HCC1806 and HCC38-CD44 Hi cell lines. *n=*3 independent experiments each with triplicate technical replicates analysed, significance measured using one-way ANOVA. (**b**) Representative images of proliferation analyses following control or SLC2A3 targeted siRNA treatment in SUM159, HCC1806 and HCC38-CD44 Hi cells. Images shown taken at experimental endpoint of 96 hours. (**c**) Representative images of tumorspheres derived from HCC1806 and HCC38-CD44 Hi cell lines following control or SLC2A3 targeted siRNA treatment. Images taken at experimental endpoint of 21 and 14 days for HCC1806 and HCC38-CD44 Hi respectively.

**Supplementary Figure 10.**
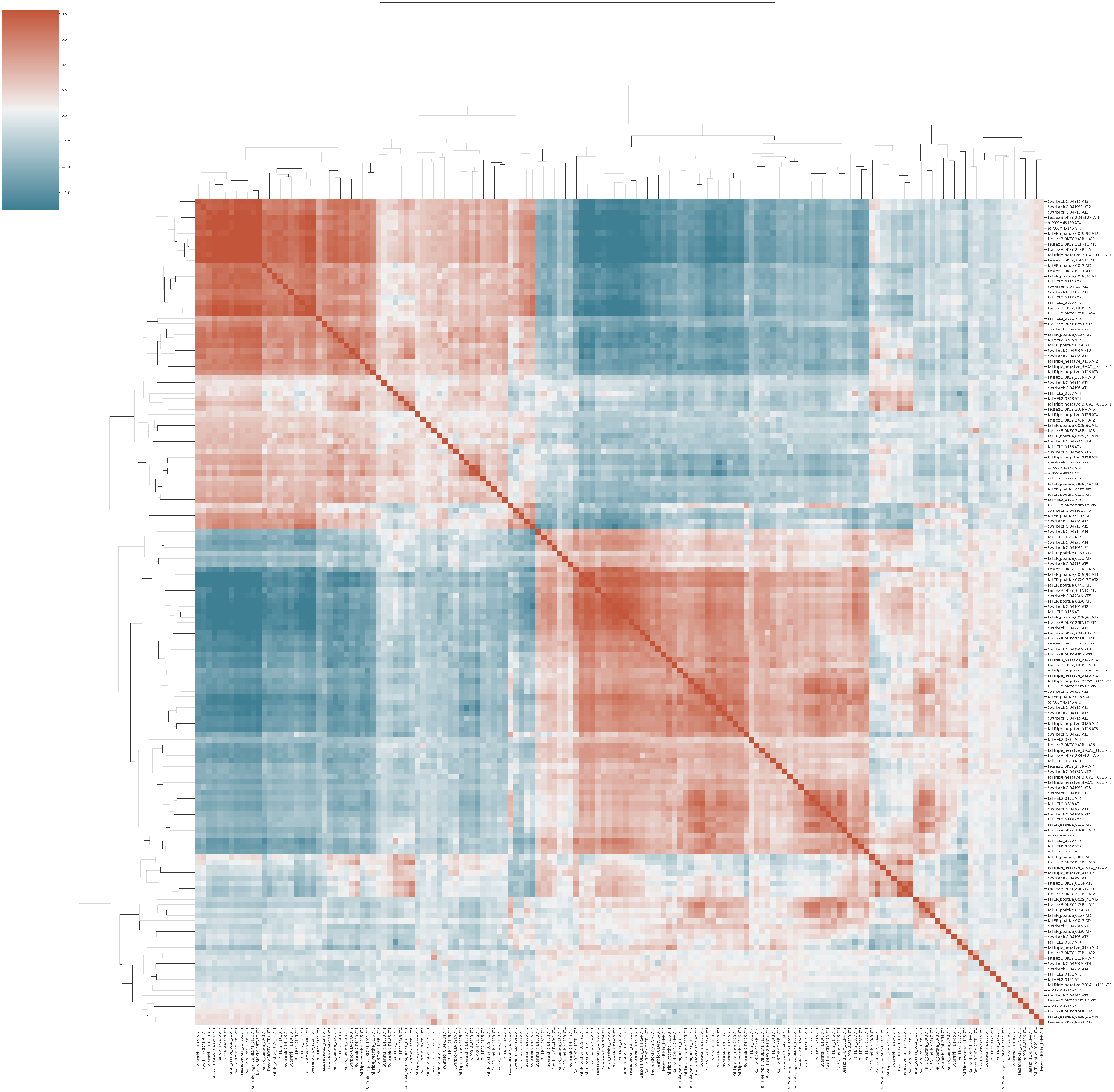
Cosine similarity across human breast cancer tumor archetypes.

## Supplemental Table Legends

**Supplementary Table 1**.

Enriched marker genes and corresponding statistics from Wilcoxon rank sum test. Sheets correspond to genes associated with each archetype identified in each Primary tumor replicate.

**Supplementary Table 2**.

Enriched marker genes and corresponding statistics from Wilcoxon rank sum test. Sheets correspond to genes associated with each archetype identified in each tissue (Primary, LN, Liver, Lung).

## Methods

### Background on cell state heterogeneity analysis

#### Clustering-based approaches unreliably identify clusters when data is a continuum of cells

Clustering is the most commonly used technique for characterizing cell state heterogeneity in single-cell data, and is considered a standard part of single-cell workflows and best practices [30]. However, clustering can be a nontrivial task, both with computational challenges and challenges with interpretation and annotation [23]. This is in part due to the fact that clustering assumes that data is composed of biologically distinct groups, such as discrete cell types.

After embedding the primary scRNA-seq data into 3-dimensions, it is evident that for the primary tumor, the cells are forming a connected manifold along the cellular state space, rather than separating into clusters (Figure 2a). After running Leiden clustering 100 times with default parameters, the cluster assignments changed at the boundaries, indicating that these cells are not strongly committed to one cluster.

We hypothesized that the cells at cluster boundaries are intermediate cells between the more distal extreme states. To test the ability of clustering-based analysis to characterize such datasets, we simulated a curved tetrahedron, where the datapoints are defined as a continuum between the vertices. We also defined a signal on the tetrahedron as a function of the affinity to one vertex (Supplementary Figure 2a). Clustering the simulated data with ten different clustering algorithms reveals (1) the lack of concordance across clustering methods when there is no latent cluster structure in the dataset and (2) the limitations of discretizing the cellular state space in characterizing continuous signals (Supplementary Figure 2b).

#### Trajectory-based approaches enforce lineage structure that do not accurately capture simulated signal

On the other hand, trajectory inference methods are commonly used to identify continuous paths in the datasets in order to define pseudotemporal ordering of cells, often for learning developmental decisions [20, 39, 42, 43]. We show that, without clear lineage structure in the dataset, trajectory-based methods are not able to learn an intelligible ordering of cells or meaningfully characterize the defined signal (Supplementary Figure 2c).

### Background on Archetypal Analysis

#### Archetypal Analysis Overview

Archetypal analysis (AA) is an unsupervised learning method that aims to find extremal points, called *archetypes*, such that every point in a dataset can be approximated as a mixture of these archetypes [11]. Given a dataset *X* = *{x*_1_, …, *x*_*N*_ *} ⊂* ℝ^*n*^, the archetypes *{z*_1_, …, *z*_*k*_*} ⊂* ℝ^*n*^ are chosen so that for each data point *x*_*i*_ there exists *α*_*i*,1_, …, *α*_*i,k*_ *∈* [0, 1] such that

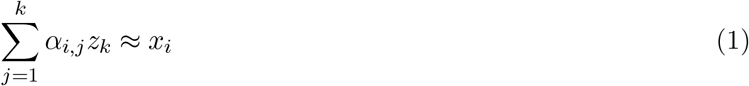

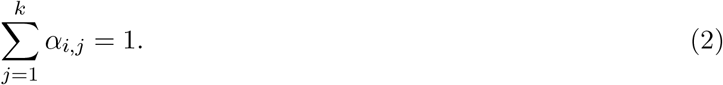

This type of linear combination where the coefficients are non-negative and sum to 1 is called a *convex combination*. The set of all such convex combinations of the archetypes *{z*_1_, …, *z*_*k*_*}* is a (*k−*1)*-simplex*. Note that the archetypes are not constrained to be points from the dataset.

#### Principal Convex Hull Analysis (PCHA)

One of the first AA algorithms was principal convex hull analysis (PCHA), proposed in [11]. PCHA constrains the archetypes to be convex combinations of the input data points. It finds these archetypes through the following optimization problem:

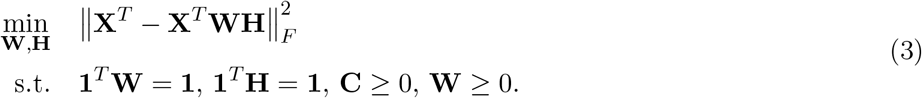

where **W** *∈* ℝ^*N×p*^ maps the data to the archetypes and **H** *∈* ℝ^*p×N*^ contains the coordinates of the archetypes in the feature space. The constraints on **W** guarantee that the archetypes are convex combinations of the data points while the constraints on **H** guarantee that the data points are convex combinations of the archetypes. [11] also defines an optimization algorithm for the expression above using alternating convex least-squares. This algorithm seeks solutions that satisfy the constraints **1**^*T*^ **W** = **1** and **1**^*T*^ **H** = **1** by adding auxiliary terms to the objective function. [33] builds on this work by modifying the optimization algorithm to use projected gradient in the alternating optimization steps to find solutions that satisfy the constraints.

#### Non-linear archetypal analysis variants (kPCHA, Javadi et al, Chen et al)

[33] also proposes kernel principal convex hull analysis (kPCHA), which is analogous to kernel principal components analysis (kPCA). PCHA is rotation equivariant, meaning that applying a rotation to the input dataset effectively results in the same rotation being applied the output features and archetypes. This implies that the output of PCHA only depends on the kernel matrix **X**^*T*^ **X** rather than the actual input dataset **X**. kPCHA takes advantage of this fact by computing this kernel matrix in a possibly-infinite dimensional reproducing kernel Hilbert space (RKHS), then running PCHA on this kernel matrix instead of **X**^*T*^ **X**. The mapping from the input feature space into the RKHS is typically non-linear, allowing kPCHA to potentially take advantage of the manifold geometry of the underlying dataset.

Several other works have also extended the algorithm proposed by [11]. In [22], the requirement that the archetypes are convex combinations of input data is relaxed. The archetypes are found by optimizing an objective function with two terms. The first term is similar to the objective function in 3 above and captures how well the convex hull of the archetypes aligns with the dataset. The second term reflects how close the archetypes are to the convex hull of the dataset. Meanwhile [9] proposes an algorithm to optimize 3 using an active-set approach.

### Background on Machine Learning

#### Autoencoders

An autoencoder is a type of neural network that is used to learn compressed representations of data. Autoencoders are comprised of two separate networks: an encoder and a decoder. The encoder network maps the input data into a low-dimensional feature space or latent space, while the decoder tries to reconstruct the original data from this low-dimensional representation. Through minimizing the error between the original data and the reconstruction, termed *reconstruction loss*, autoencoders have been shown to successfully learn the structure of data, and have had particular utility in capturing a meaningful representation of single-cell data [3, 14, 19, 29, 46].

#### Manifold learning

Manifold learning is a subfield of machine learning built around the manifold hypothesis, which asserts that high-dimensional datasets are sampled from low-dimensional manifolds that lie in the high-dimensional space. Here a *manifold* refers to a space that is locally isomorphic to a Euclidean space. Many methods in unsupervised learning attempt to implicitly or explicitly capture the structure of the underlying data manifold. For instance, the latent space of an autoencoder can be viewed as a parameterization of the data manifold.

#### AAnet Overview

##### AAnet architecture

AAnet is designed to have a flexible number and size of layers depending on the complexity for the task. For our purposes, we found that a 2-layer (256 nodes, 128 nodes) encoder and (128 nodes, 256 nodes) 2-layer decoder worked well. The batch size was 256, the optimizer was ADAM, the learning rate was set to 1e-3, and the weight initialization was Xavier. All hidden layers contain Tanh nonlinear activations, besides layers directly before and after archetypal layer which are linear so that each point is a linear combination of archetypes. The default weight on *extrema loss γ*_*extrema*_ is set to 1. To encourage the archetypes to be tight, i.e. close to the data, we can add Gaussian noise *∼ N* (0, 0.05) in the latent layer during training. For all datasets, we reduced dimensionality using PCA before running AAnet and inverse-transformed the learned archetypes to the ambient space.

##### Reconstruction Loss

The main loss function for the autoencoder seeks to minimize the difference between the original input fed into the encoder *z* = *E*(*x*) and the reconstructed input produced by the decoder 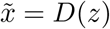, termed *reconstruction loss*. Standardly, autoencoders use the mean squared difference of these two terms:

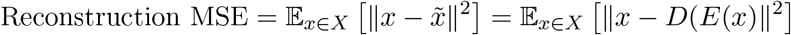

##### Archetypal Loss

In addition to the reconstruction loss, we want to enforce the latent space of the autoencoder to learn the structure of the data with respect to the archetypes. To this end, we convert the coordinates from Cartesian to barycentric after the encoder learns the transformation. The barycentric coordinate system, related to Cartesian coordinates, is a system in which each point is specified by reference to a simplex. When coordinates are normalized to sum to 1, the vertices of the simplex are denoted by *k+1* one-hot vectors of length *k+1* for a *k*-simplex. For example, a triangle is a 2-simplex with 3 vertices, where the 3 vertices are (1,0,0), (0,1,0), and (0,0,1). All coefficients of point *P* are positive if and only if *P* is inside the simplex.

As this coordinate system describes points with respect to a *k*-simplex, it is well-suited to be the latent space for *k* archetypes.

To enable interpretation of points as convex combinations of archetypes, we enforce each point stays within the simplex by adding an *archetypal loss* term, the mean squared error of the negative coefficients:

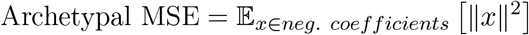

##### Extrema Loss

We developed a novel method to identify *k* plausible archetypes prior to model training. This method, explained in detail below, builds a graph from the data and then uses the eigenvectors of the Laplacian matrix to find extreme points in the datasets; these points will be refered to as *diffusion extrema*. We then include an *extrema loss* term that penalizes large distances in the latent space between the diffusion extrema and the vertices of the simplex. If the diffusion extrema and standard basis vectors in the latent space ℝ^*k*^ are labelled as 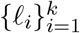 and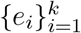, respectively, then this loss term can be calculated as

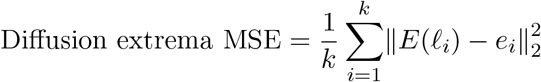

Let *{x*_1_, …, *x*_*n*_*} ∈* ℝ^*m*^ be the points in the dataset. Then the procedure for finding these diffusion extrema is as follows:

1. Construct a graph *G* from the dataset. This can be done by computing a symmetrized *k*-nearest neighbors graph from the dataset and then weighting the edges with a Gaussian kernel, as is done in [32].
2. Let ***ψ***_1_, …, ***ψ***_*n*_ denote the eigenvectors of the combinatorial Laplacian ***L*** of *G* with corresponding eigenvalues *λ*_1_ *≤ … ≤ λ*_*n*_. Compute

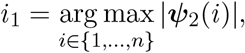

where ***ψ***_2_(*i*) is the *i*th entry of ***ψ***_2_.
3. Let 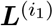 denote ***L*** with the entries in the *i*th row and *i*th column replaced by zeros. Likewise let 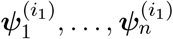 denote the eigenvectors of 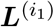, again ordered in an ascending fashion by corresponding eigenvalue. Compute

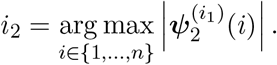
4. Let 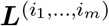 denote ***L*** with the entries in the *i*_1_th, …, *i*_*m−*1_st, and *i*_*m*_th rows and columns replaced by zeros. Likewise let 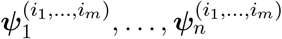 denote the eigenvectors of 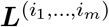. Iteratively for each *j* = 3, …, *k* compute

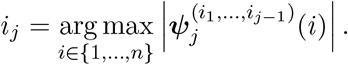
5. The diffusion extrema are 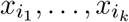.

The intuition behind this algorithm comes from an application of Courant-Fischer theorem for symmetric matrices. Given an *n × n* symmetric matrix ***A*** with eigenvectors ***a***_1_, …, ***a***_*k*_, Courant-Fischer tells us that

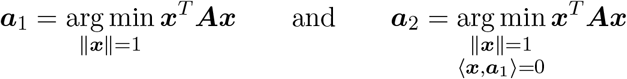

If ***A*** is the Laplacian matrix ***L*** for some weighted graph *G* = (*V, E, w*) then

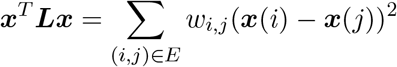

Intuitively the quadratic form ***x***^*T*^ ***Lx*** captures how smoothly **x** varies over the edges of *G*. Hence ***ψ***_1_ is the (normalized) constant vector, while ***ψ***_1_ can be viewed as a smooth signal on *G* that is orthogonal to the constant vector. Now if we consider the matrix ***L***^(*i*)^ we see that minimizing the quadratic form ***x***^*T*^ ***L***^(*i*)^***x*** can be recast as minimizing ***x***^*T*^ ***Lx*** with the additional constraint that ***x***(*i*) = 0, i.e.

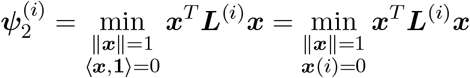

This is because 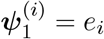, the *i*th standard basis vector. Extending this reasoning to 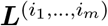, where *i*_1_, …, *i*_*m*_ *∈ {*1, …, |*V* |*}* are unique but arbitrary, we see that 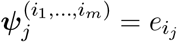 for 1 *≤ j ≤ m*. Hence

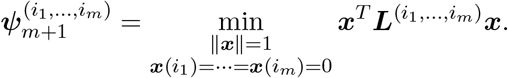

This tells us that the *m*th eigenvector of 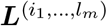 is a signal on *G* that is as smooth as possible while also having the value 0 on vertices *i*_1_, …, *i*_*m*_. Then we should expect 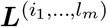 to attain its largest absolute value at a vertex that is far from the vertices *i*_1_, …, *i*_*m*_. This is the guiding principle behind the above diffusion extrema method. Step (2) takes advantage of the fact that ***ψ***_2_ is smooth on *G* and therefore is likely to attain its maximum absolutely value at extremal points in the graph. The vertex at which ***ψ***_2_ has its largest absolute value is chosen to be the first diffusion extrema. Steps (3) and (4) then use the properties of 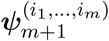 discussed above to iteratively find vertices in the graph that are far away from the extrema that have already been chosen.

It has not been conclusively proven that the eigenvectors of the Laplacian ***L*** and Laplacian variant 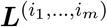 attain their maximum/minimal values at extremal points in the graph, nor is it entirely clear which vertices in a graph can be called in extremal. However through our experiments on both toy data and real biological data we believe this diffusion extrema method to be robust and effective in identify reasonable archetypes in a dataset.

#### Choosing number of archetypes

To choose the number of archetypes, we first define a range of possible archetype counts *m* = 2, 3, …, *n*_*at*_. Then, we calculate *n*_*at*_ diffusion extrema. For the *m*^*th*^ diffusion extremum, we calculate the geodesic distance of that extremum to all previous extremum (first to *m −* 1^*th*^) and take the mean of all the distances. Intuitively, this score tells how transcriptionally distinct archetype *m* is from all existing archetypes, and if it is very similar, it is likely not a new archetype. Thus, we take the knee point of this score as the number of archetypes *k* for downstream analysis.

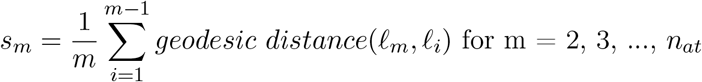

### Comparative Analysis

#### Simulated Datasets

To generate a simulated dataset for comparisons, we sample non-uniformly from a k-simplex projected onto a k-dimensional sphere using a stereographic projection. This enables comparisons on a dataset with nonlinear geometry and known vertices.

For comparisons in Supplementary Figure 1, we sample 10,000 points from a 3-simplex (a tetrahedron) projected onto a hypersphere. This results in a curved tetrahedron, where the ground truth archetypes do not correspond to the extrema in the data space (as they would for a classic tetrahedron). We embed the curved tetrahedron with PHATE [32], a dimensionality reduction tool designed to capture nonlinear local and global variation.

Next, we simulated a signal, defined by the *sine* of the affinity between vertex 4 and all other points in the tetrahedron based on data geometry. The ground truth signal is thus a sinusoidal function. This signal is defined by its relation to a ground truth archetype, and is designed to model biological processes that are enriched with respect to a cellular archetype.

### Comparison to clustering methods

To compare archetypal analysis to clustering on the curved tetrahedron, we clustered the data using default parameters for 10 different clustering algorithms in sklearn.cluster. Five methods required the number of clusters to be specified, and for these we specified four clusters, the ground truth number of vertices for a tetrahedron.

We ran clustering using ten different clustering algorithms: five that require the number of clusters to be specified (KMeans, Agglomerative Clustering (Ward), Agglomerative Clustering (Single Linkage), Agglomerative Clustering (Complete Linkage), and Spectral Clustering), and five that infer the number of clusters from the data (Affinity Propagation, Mean Shift, DBSCAN, OPTICS, and BIRCH).

### Comparison to trajectory-inference methods

For trajectory inference comparisons, we ran Slingshot and diffusion pseudotime (DPT). Slingshot requires cluster labels and a dimensionality-reduced representation. We used the PHATE embedding and the KMeans cluster labels as input for Figure 2, as KMeans is a popular method for clustering that produced stable results. Besides these two inputs, all other parameters were default. DPT was also run with default parameters, with the required starting cell chosen randomly from cluster 3 of KMeans (the cluster containing extremum 4).

### Comparison to archetypal analysis methods

All archetypal analysis methods require input of the number of archetypes, so we specified four archetypes for each method, otherwise running with default parameters. To generate MSE calculations between the ground truth vertices and inferred archetypes, we ran each method 5 times and visualized the first run for each method.

By the definition of barycentric coordinates, a point has a value closer to 1 for dimension *k* if it has a high affinity to the *k*-th vertex, and a value closer to 0 if it has a low affinity to the *k*-th vertex. Therefore, we color the dataset with the latent coordinates for each dimension to determine if the latent space has semantic structure, and if AAnet is effectively learning affinity relative to each extremum.

### AAnet for published antigen-specific CD8+ T cells

For each dataset, we ran TruncatedSVD to reduce the dimensionality to 100, and then ran AAnet with default parameters. The expression levels in Supplementary Figure 2 were z-score normalized and archetypes were clustered based on cosine similarity.

### Computational Methods for TNBC

#### Single-cell RNA preprocessing

The CellRanger Analysis Pipeline (v3.0.2) was used to align the sequencing reads (fastq) to a pre-built reference genome (10x Genomics) containing both the human and mouse genomes (GRCh38 + mm10) and gene expression quantified in each cell. Cells from four primary tumors were sequenced (T1-T4) with a total of 33,938 cells sequenced (T1 = 8707, T2 = 5951, T3 = 8934, T4 = 10346). Gene expression data was loaded into Python (v3.8) and quality-control (QC) statistics were computed using the scanpy.pp.calculate_qc_metrics function (v1.9.1, [49]). To ensure only human tumor cells were taken for downstream archetypal analysis, cells with less than 99% of data aligning to the human genome were removed. Mouse genes were also removed prior to calculating library size. Cells with a total library size between 2000-50,000 UMI and expressing at least 1000 genes were retained for downstream analysis. A total of 28,478 primary tumor cells passed QC filtering (T1 = 7606, T2 = 5118, T3 = 8163, T4 = 7591). Remaining tumor cells were normalized to 10,000 reads per cell and square-root transformed using the scprep package (v1.2.3, github.com/krishnaswamylab/scprep). To correct for dropout data were smoothed with the manifold smoothing method MAGIC (v3.0.0, [45]). Cell numbers remaining after QC were visualized using the R package ggplot2 (v3.4.2, [48]). Cell-cycle phase assignment was performed using the `scanpy.tl.score_genes_cell_cycle` function with previously defined S-phase and G2M-phase gene lists [41].

Highly variable genes were detected within each sample using the scprep function `scprep.select.highly_variable_genes` and a cellular graph constructed based on KNN and alpha decay kernel using graphtools (v1.5.3, github.com/krishnaswamylab/graphtools). For the combined analysis of all tumors we used an MNN kernel to build a cellular graph with batch correction between the replicates. We then used MAGIC (v3.0.0, [45]) to transform each graph into the gene space, and ran TruncatedSVD to reduce the dimensionality to 100 and visualize samples in reduced dimensions.

### AT similarity, affinity and commitment

AAnet was run on both individual samples and a combined dataset using default parameters. Data archetypes were defined for each dataset using AAnet and archetypal expression vectors were generating by transforming archetype coordinates back into the gene space. Cosine similarity was calculated between expression vectors and hierarchical clustering used to identify five orthologous expression states between datasets. The affinity of each cell to each archetype was calculated based on the distance of each cell to each archetype in the AAnet latent embedding. As the combined affinity of each cell to all archetypes is regularized to 1, cells with an affinity to single archetype greater than the sum of their affinity to all other archetypes (i.e. with an affinity >0.5 for any archetypes) were defined as committed to that archetype. The archetypal composition of each sample was determined by summing the number of committed cells per archetype and assigning cells with affinity for all archetypes <0.5 as uncommitted.

### Biological characterization of archetypal states

Marker genes upregulated in each archetype were calculated by comparing gene expression between cells that were the most committed to each archetype. The top 125 cells with the strongest affinity for each archetype in the combined AAnet latent space were selected from each replicate (500 cells per archetype), as well as the 125 cell per replicate furthest from any archetype as the uncommitted populations (500 cells uncommitted). Genes that were upregulated in any group relative to all other groups (FDR < 0.05) were determined using the FindMarkers function from the Seurat R package (v4.0, [40]). Cancer hallmark gene sets [28] overrepresented (FDR < 0.1) in markers associated with each archetype were determined by 1-sided Fisher’s exact test using the clusterProfiler R-package (v3.16.1, [52]) and visualized using ggplot2.

### Spatial RNA data generation

Spatial transcriptomics was conducted using 10X Visium Spatial Gene Expression Slide and Reagent Kit, 16 rxns (PN-1000184), according to the protocol detailed in document CG000239RevD for the TNBCs and CG000239RevE for the Xenografts, available in 10X Genomics demonstrated protocols. Cryo-sectioning was done on OCT embedded and snap frozen tissue samples at 10um thickness and placed on cold Visium slide arrays. The sections were adhered by swiftly warming of the backside of the slide. The slides were then kept in −80°C less than 4 weeks before processing accordingly.

In short, the slides were first warmed at 37°C for one minute and then immersed in pre-chilled methanol (VWR EU, 20847.307) for 30 minutes at −20°C for fixation. Staining with hematoxylin and eosin was carried out by one min of drying of the tissues with 500uL of isopropanol (Fisher Scientific, A461-1) followed by air drying until sections turned white. Around 1 mL of with Mayer’s hematoxylin (Agilent, S23309) was pipetted onto the slides and treated for four minutes. The slides were then washed in nuclease free water followed by a two-minute incubation with bluing buffer (Agilent, CS702), washed again and then counterstained with buffered eosin (Sigma-Aldrich, HT110216, 1:10 dilution in Tris-Acetic Acid Buffer). The slides were air dried for about 2 minutes and then warmed for 5 minutes at 37°C before mounting using 85percent Glycerol (Merck, 104094) and a coverslip.

Bright field histology images were obtained using a 20X objective on Zeiss microscope using the Metafer VSlide system and the images were processed by the VSlide software. The images were extracted as jpgs for downstream analysis.

After imaging, the coverslip and the remaining glycerol was washed off in Milli-Q water and the slides were attached in plastic cassettes included in the reagent kit and first subjected to 20 minutes of permeabilization at 37°C to let the mRNA reach the probes on the slide surface for binding.

The protocol was then followed without deviations to create amplified libraries which in the end were individually indexed using the Dual Index Kit TT SetA, (PN-1000215, 10X Genomics), quality controlled on a BioAnalyzer instrument and concentrations were measured using Qubit DNA HS. The libraries were pooled equimolarly (2nM) and sequenced on the Nextseq 500 (Illumina platform) for the tnbcs and Nextseq 2000 (Illumina platform) for the xenografts. To reach the appropriate read depth the recommended number of reads per ST spot were applied according to the protocol.

### Spatial RNA preprocessing

The SpaceRanger Analysis Pipeline (v2.0.0, 10x Genomics) was used to align the sequencing reads (fastq) to a pre-built reference genome (10x Genomics) containing both the human and mouse genomes (GRCh38 + mm10) and gene expression quantified in each cell. Quality-control (QC) statistics were computed using the STutility package [7]. Voxels with a library size < 3000 UMI were removed and remaining voxels manually curated to remove those that were disconnected from the main tissue section. Human and mouse genes were separated to create two datasets per sample, one measuring the expression of human genes at each voxel, pertaining to tumor cells, and the other measuring expression of mouse genes, pertaining to cells from the microenvironment. Genes were removed if they were detected in less than 10 voxels. Finally each dataset was filtered to remove voxels with low library diversity for a given genome (<1000 human genes detected, <300 mouse genes detected). After QC filtering, the human dataset comprised of 14,819 genes measured at 1170 and 1105 voxels for each tumor, while the mouse dataset comprised of 12,119 genes measured at 1150 and 1079 voxels in TX and TX section respectively. Remaining datasets were each normalized to 10,000 reads per voxel and log transformed. To correct for dropout (human dataset = 69.8% zeros, mouse dataset = 90.6% zeros) data were smoothed with the manifold smoothing method MAGIC. Cell-cycle phase assignment was performed using the CellCycleScoring function in Seurat [40] with previously defined S-phase and G2M-phase gene lists [41]. Spatial plots throughout the manuscript were generated using STUtility.

### Mapping of archetypes to spatial transcriptomics with scMMGAN

To map archetypes identified in the single-cell data from primary tumors to voxels in the spatial transcriptomic samples we used scMMGAN [4]. Smoothed expression data were zero-centered and unit scaled each dimension, and reduce to 50 principal components used as input to the scMMGAN generator, with the combined scRNAseq data described above used as input for the discriminator layer. scMMGAN was run with a generator consisting of three internal layers of 128, 256, and 512 neurons with batch norm and leaky rectified linear unit activations after each layer, and a discriminator consisting of three internal layers with 1,024, 512, and 256 neurons with the same batch norm and activations except with minibatching after the first layer. We use the geometry-preserving correspondence loss with a coefficient of 10, cycle-loss coefficient of 1, learning rate of 0.0001, and batch size of 256. This network was used to generate a single-cell-like representation of each spatial voxel. These generated single-cell values were then embedded into the AAnet latent space trained using the combined single-cell RNAseq dataset. The affinity of each voxel to each archetype was then determined based on the distance of its generated single-cell values to each archetype in the trained AAnet latent space. Each voxel was then assigned to an archetype to which it had the highest affinity, or uncommitted in the case the maximum affinity corresponded to more than one archetype. It is important to note that the resolution of spatial transcriptomics voxels above the single-cell level (estimated between 3-10 cells per voxel), thus archetypes represent the dominant expression state among cells in the voxel. We also scored voxels based on the expression of the marker gene sets associated with each archetype in the single-cell data using on the first principal component of gene set expression, analogous to the eigengene metric used to summarize gene coexpression networks [25]. Scores were calculated based on expression of the top 50 marker genes with the highest log fold enrichment per archetype, excluding mitochondrial and ribosomal genes.

### Microenvironment mapping and enrichment

We used data aligning to the mouse genome at each voxel to analyze the microenvironment associated with each archetype. Based on the archetypal assignment of voxels using tumor data, as described above, we identified differentially expressed genes between microenvironment spatially colocalized with each archetype using the FindMarkers function in Seurat (LFC >0.1, FDR<0.05) [40]. Enrichment of Gene Ontology Biological Processes and cell-types markers associated with “Connective tissue”, “Immune system”, “Smooth muscle”, “Epithelium”, “Vasculature”, “Blood”, “Mammary gland” or “Skeletal muscle” in the Pangloa database (v27/03/2020) [18] among differentially expressed genes was determined with a 1-sided Fisher’s exact test using the clusterProfiler R-package [54]. Putative ligand-receptor interactions between archetypes were identified using CellPhoneDB (v2) [13]. Human orthologs to mouse genes were idenitfied in biomaRt and counts matrices were merged based on gene id prior to running CellPhoneDB using parameters –iterations 1000 –threshold 0.2. and identifying significant interactions (FDR < 0.05). The metabolism between archetypes and microenvrionment was compared based expression of key enzymes involved in glycolysis and the tricarboxylic acid (TCA) cycle in each archetype and associated microenvironment. Glycolytic enzymes were designated as either hypoxic based on their enriched expression (LFC > 0) in voxels assigned to the hypoxic AT5 archtype. Heatmaps were generated based on the mean of scaled values for voxels associated with each archetype (human genes) or microenvironment (orthologous mouse genes).

### Cell Culture

SUM159 cells were cultured in Ham’s F12 + 1% (v/v) hydrocortisone, HCC1806 and HCC38-CD44 Hi cells were cultured in RPMI media. Media were supplemented with 10% (v/v) fetal bovine serum (GE healthcare) and 1% (v/v) pencillin-streptomycin. To isolate CD44Hi cells from the parental HCC38 cell line, cells were expanded *in vitro* in 150 mm tissue culture-treated culture dish (Corning). For the initial FACS rounds, cells were trypsinized and 1-2 x 10^7^ cells were stained for the membrane marker CD44 (BD anti-human CD44-PE-cy7 (1:800)) for 25 min at 4C. Antibody titration was previously determined staining a battery of breast cancer cell lines. Pure CD44^Hi^ cells were collected and replated for expansion in culture. Following sorting purification, cultures were supplemented with 0.1% (v/v) gentamicin and 1% (v/v) antibiotic-antimycotic for 2 passages to avoid contamination. Sequential rounds of FACS enrichment were performed until 100% pure populations were isolated. Data collection was performed using a BD Aria III and FACSDiva software (BD Biosciences). Flowjo X10.7.1 was used for data analysis.

#### siRNA treatment

Cells were plated, allowed to grow for 24h then transfected with either 20nM siRNA *Silencer* ™ Select negative control (Invitrogen, 4390843) or SLC2A3 directed siRNA (Invitrogen, 4390824). siRNAs were transfected using Optimem Reduced Serum Media (ThermoFisher 31985070) and Lipofectamine 3000 (ThermoFisher scientific L3000015#) as per standard protocol. Cells were harvested at 24h for qPCR analysis or trypsinized and re-seeded for proliferation and/or tumorsphere assay.

#### qRT-PCR

Total RNA was extracted and purified using the RNeasy micro kit (Qiagen). Reverse transcription was performed from 1 μg of total RNA with Superscript IV reverse transcriptase (Life Technologies, CA, USA) according to the manufacturer’s instructions. The reverse transcription product was diluted 1:10 with TaqMan Fast Advanced Master Mix (Invitrogen, 4444556) and used as a cDNA template for qPCR analysis. Real-time quantitative PCR was performed using the QuantStudio™ 7 Flex Real-Time PCR System (Applied Biosystems, CA, USA). Results are represented as mean values normalised to controls.

#### Proliferation

Following 24 hours of treatment of control or SLC2A3 targeted siRNA treatment, cells were seeded (1000 per well) in a 96-well plate. Proliferation was determined by confluence (%) per well, measured every 24 hours over a 96-hour using an Incucyte (Sartorius).

#### Tumorsphere-forming assay

After 24 hours of treatment of control or SLC2A3 targeted siRNA treatment, cells were trypsinized and seeded in an ultra-low attachment 96 well plate (Corning) (300 cells per well) at 1000 cells per well. Tumorsphere media was comprised of methylcellulose (Sigma, m-7027) and basal media supplemented with 20ng/ml basic fibroblast growth factor (Millipore, GF003), 20ng/ml human epidermal growth factor (Sigma, E1264), B27 (Life Technologies, 17504-044) and 4μg/ml heparin (Sigma, H3149). 50ul additional tumorsphere media was added every 5 days and tumorspheres were counted and imaged at 14 or 21 days.

#### Human scRNA-seq data preprocessing

For experiments from [6, 35, 51], the datasets were first subset to cancer epithelial cells based on prior annotation, and only samples with *≥* 1000 cells were analyzed with AAnet. We then further preprocessed the data by removing genes expressed in fewer than 5 cells, filtering library size to between 1500 and 60000 UMI counts, and L1 normalizing for library size. We then square-root transformed the data, and filtered any remaining contaminating cells based on marker gene expression. Finally, we embedded the data with MAGIC and PCA before identifying archetypes with AAnet. For experiments based on the PDX model, the same pipeline was followed, where additionally cells with less than 99% of reads aligning to human cells were removed, and mouse genes were removed.

## Data Availability

The accession codes to the newly generated single-cell and spatial data will be provided before publication. Public CD8+ T cell data can be accessed at GEO under accession number GSE182509. Processed scRNA-seq data from [35] is available as GEO series GSE161529. Raw sequencing reads of all single-cell experiments from [6] have been deposited in the European Genome-phenome Archive (EGA) under study no. EGAS00001004809. Processed scRNA-seq data from [51] are also available through the Gene Expression Omnibus under accession number GSE176078.

## Code Availability

The source code is available at https://github.com/KrishnaswamyLab/AAnet.

## Author Contributions

A.V., D.B.B., A.B., D.V.D., and S.K. developed AAnet. C.L.C. and B.P.S.J. designed the biological experiment. B.P.S.J. performed biological experiments. M.A. aligned scRNA-seq and spatial transcriptomic data with scMMGAN. S.E.Y., A.V., and C.P. performed computational analyses. A.M. and J.K. performed and supported spatial transcriptomic experiments. J.H. performed analysis and interpretation of metabolomic data. L.D.G. and S.K. oversaw computational analyses done by S.E.Y., A.V., and C.P. A.V., S.E.Y., S.K. and C.L.C. wrote the manuscript.

